# Deciphering the developmental order and microstructural patterns of early white matter pathways in a diffusion MRI based fetal brain atlas

**DOI:** 10.1101/2022.02.28.481882

**Authors:** Ruike Chen, Cong Sun, Tingting Liu, Yuhao Liao, Junyan Wang, Yi Sun, Yi Zhang, Guangbin Wang, Dan Wu

## Abstract

White matter of the fetal brain undergoes rapid development to form the early structural connections. Diffusion MRI has shown to be a useful tool to depict the fetal brain white matter *in utero*, and many studies have observed increasing fractional anisotropy and decreasing mean diffusivity in the fetal brain during the second-to-third trimester, whereas others reported non-monotonic changes. Unbiased diffusion MRI atlases of the fetal brain are important for characterizing developmental trajectories of white matter and providing normative references for in-utero diagnosis of prenatal abnormalities. To date, the sole fetal brain diffusion MRI atlas was collected from a Caucasian/mixed population, and was constructed based on the diffusion tensor model with limited spatial resolution.

In this work, we proposed a fiber orientation distribution (FOD) based pipeline for the generation of fetal brain diffusion MRI atlases, which showed better registration accuracy than diffusion tensor-based pipeline. Based on the FOD pipeline, we constructed the first Chinese fetal brain diffusion MRI atlas using 89 normal fetal diffusion MRI scans at GA between 24 and 38 weeks. Complex non-monotonic trends of tensor- and FOD-derived microstructural parameters in eight white matter tracts were observed, which jointly pointed to different phases of microstructural development. Specifically, we speculated that the turning point of the diffusivity trajectory may correspond to the starting point of pre-myelination, based on which, the developmental order of white matter tracts can be mapped and the order was in agreement with order of myelination from histological studies. The normative atlas also provided a reference for detection of abnormal white matter development, such as that in congenital heart disease. Therefore, the high-order fetal brain diffusion MRI atlas established in this study depicted the spatiotemporal pattern of early white matter development, and findings may help decipher the distinct microstructural events *in utero*.

## Introduction

White matter (WM) tracts, composed of axonal bundles, are the pathways in the brain for transmitting information between different cortical and subcortical regions in the brain. During the second to third trimester of gestation, the WM fibers undergo several developmental stages including the fasciculation of axons, oligodendroglial proliferation and differentiation, formation of pre-myelin sheaths around the axons in preparation of myelinogenesis, and true myelination process.^1–3^ The detailed processes in the development of WM microstructure, such as myelination, has been explored mainly by histological studies in postmortem fetal brains.^4–8^ However, fetal brain specimens are difficult to obtain, and the traditional histological methods are limited to cross-sectional slices, which lack the spatiotemporal resolution for global/intact and continuous mapping of fetal brain development. Moreover, the postmortem brains are likely to differ from the normal brains in utero. Therefore, in-utero approaches may provide more valuable information about WM development in this critical period.

Diffusion magnetic resonance imaging (dMRI) is the most useful neuroimaging tool to depict WM microstructures *in vivo*, by probing restricted water diffusion in biological tissues. For instance, diffusion tensor imaging (DTI) is most widely used to model the 3D anisotropic water diffusion in WM tracts, based on which, fractional anisotropy (FA), axial diffusivity (AD), radial diffusivity (RD) and mean diffusivity (MD) metrics are derived to reflect the microstructural integrity of WM.^9,10^ However, it only models the primary fiber direction and is limited in interpreting complex WM organization or providing specific microstructural information. High-order models such as constrained spherical deconvolution (CSD) estimates a fiber orientation distribution (FOD) function based on dMRI signals to resolve multiple fiber components, enabling fiber tractography to estimate the crossing WM fiber bundles of the brain.^11^ dMRI-based studies of the brain WM have been extensively presented in the adult, aging, and diseased brains, yet, dMRI of the perinatal brain has been difficult, primarily due to the severe motion and lack of resolution. With the recent advance of fast imaging and post-processing methods, feasibility of dMRI to image the early brain development has been demonstrated.^12–14^ For example, DTI has been used to characterize the developing WM in the infants.^15,16^

In recent years, DTI has been attempted in the fetal brain in-utero. Many of existing studies reported a significant linear increase of FA and decrease of MD or apparent diffusion coefficient (ADC) over GA.^17–22^ Others, however, observed an initial increase of ADC or non-linear changes of DTI measures in some WM areas.^19,20,23–27^ Schneider, et al.^19^ studied 78 fetuses of GA from 23 to 38 gestational weeks and spotted an initial increase of ADC values in supratentorial areas, followed by a decrease after the 30 weeks of GA. The same group conducted a longitudinal study on 28 fetuses at 21 to 34 weeks and observed an insignificant increase of ADC in frontal WM.^20^ Similar trends of ADC in frontal WM were also reported by Hoffmann, et al.^23^ and Korostyshevskaya, et al.^24^ Zanin, et al.^25^ studied WM tracts of 17 fetuses at GA between 23 and 38 weeks, whose DTI parameters were fitted to third order polynomial functions. These inconsistencies were partially related to the limited sample size, insufficient GA range, study population, and data acquisition and processing strategies in the previous studies. Also, the existing studies mostly focused on the DTI model without taking advantage of advanced dMRI models to investigate the microstructural details.

Beyond the study of individual fetal brains, spatiotemporal atlases of the fetal brain, which represent the population averaged template of fetal brain at each stage, are desired to provide normative templates and severe as reference for in-utero diagnosis of abnormal development. To the best of our knowledge, the only publically available in-utero fetal brain dMRI atlas is the CRL fetal brain DTI atlas constructed using 67 dMRI scans of normal fetuses at GA between 21 and 39 weeks.^28^ However, the CRL atlas was reconstructed with diffusion tensor model as the data were acquired from only 12 diffusion directions, and thus lack higher order descriptions of complex fiber structures. Moreover, the inter-subject registration in the CRL atlas generation pipeline was not optimized, resulting in relatively blurred population averages. Wilson, et al.^26^ studied 113 fetuses at GA from 22 to 37 weeks using weekly population-based FOD templates (not publically available), but did not further characterize the templates. The existing dMRI atlases were both collected from the Caucasian or mixed populations, while it known that there are significant brain structural differences between Chinese and Caucasian populations in adults and children.^29–32^ It has also been reported that there were anatomical differences in several neural structures between Chinese and other ethnics revealed by MRI and optical coherence tomography.^33,34^ It would be interesting to see whether these differences begin at the very beginning in the fetal brain when there is minimal environmental influence.

Therefore, the goals of the current work were: 1) to develop an optimized dMRI pipeline for accurate generation of fetal brain dMRI atlas with high-order analysis; 2) to establish a 4D spatiotemporal dMRI atlas of the Chinese fetal brain; and 3) to decipher the pattern and order of WM development with multi-modal dMRI parameters. We hypothesized the WM development during second-to-third gestation can be characterized into several phases according to the distinct developmental trajectories of the dMRI features that correspond to the ongoing microstructural events in the fetal brain. We further applied the normative atlas for analysis of abnormal fetal brain development, and also compared the proposed Chinese fetal brain dMRI atlas with CRL atlas.

## Materials and methods

### Subjects

135 pregnant women were enrolled at Shandong Provincial Hospital under the approval of the Institutional Review Board. All subjects provided written informed consent. The enrollment criteria included: (1) gestational age (GA) between 20-40 weeks; (2) no maternal comorbidities; and (3) fetuses with no clinical or ultrasound or MRI evidence of brain abnormalities. In addition, twenty-four fetuses diagnosed with congenital heart disease (CHD) through clinical ultrasound examination were enrolled in this study. The inclusion criterion for CHD was fetal echocardiogram confirmation of complex CHD according to established guidelines ISUOG.^35^ Exclusion criteria for both CHD and the normal group were as follows: (a) pregnant women with gestational diabetes mellitus or hypertensive disorder complicating pregnancy (b) multiple pregnancies; and (c) fetuses with chromosomal or genetic abnormalities diagnosed by amniocentesis.

### Data acquisition

All scans were performed on a 3T MR scanner (MAGNETOM Skyra, Siemens Healthcare, Erlangen, Germany) using a diffusion-weighted echo-planar imaging sequence with a b-value of 600 s/mm^2^, 30 gradient directions, eight non-diffusion-weighted images, 1.73 mm in-plane resolution in axial slices, 4 mm slice thickness, and two averages. Other acquisition parameters were: field of view (FOV) = 260×260 mm2, repetition time (TR) = 3900 ms, echo time (TE) = 80 ms. We excluded scans with severe motion, noise or other quality concerns, which resulted in 89 remaining scans acquired at gestational age (GA) from 24 to 38 weeks, with five to eight subjects in each week (Figure 1). Eleven of the twenty-four CHD fetuses were selected after excluding data with bad quality, with GA ranging between 25 and 30 weeks. Sex of these fetuses were unavailable due to the privacy policy.

**Figure 1.**
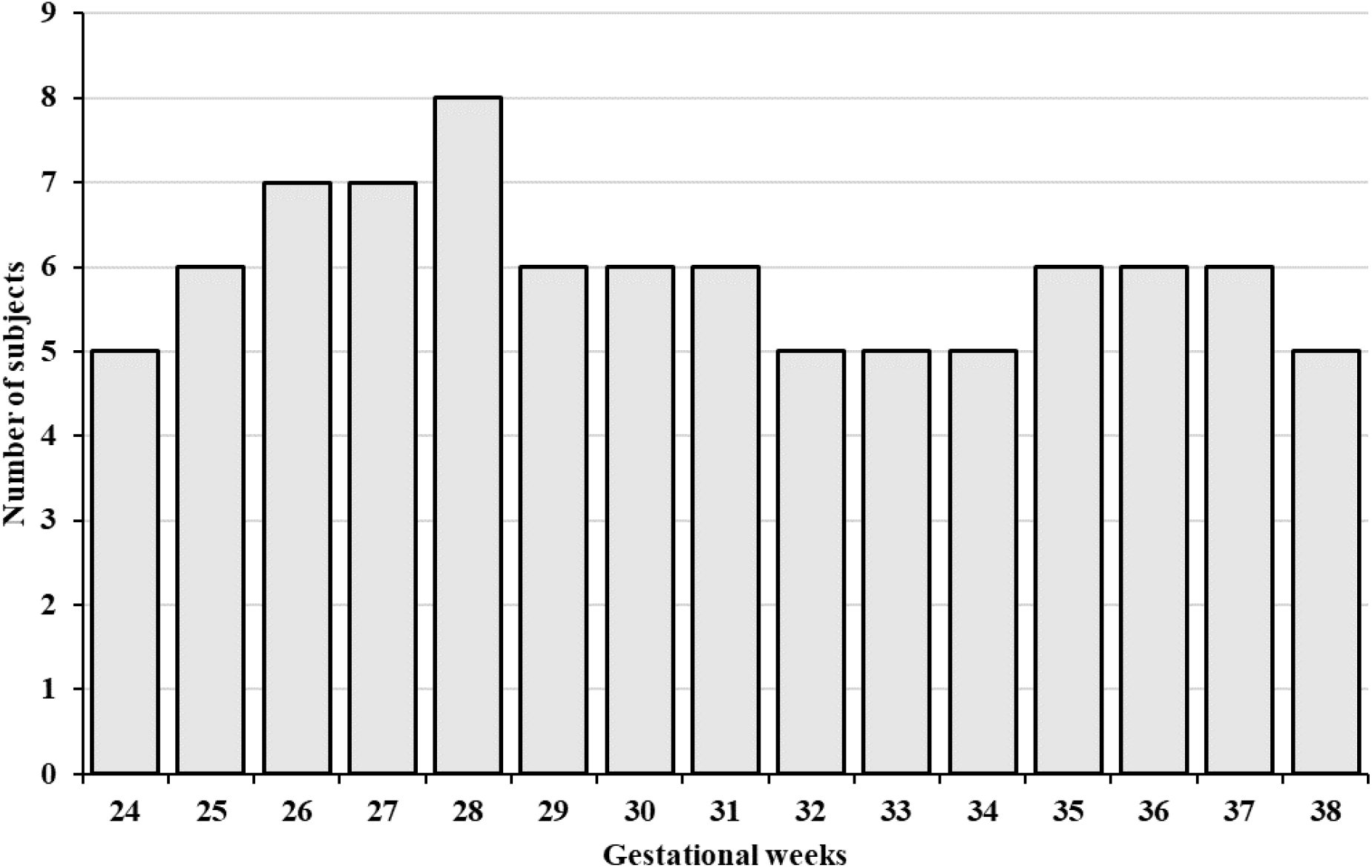
Number of normal fetal brains used in each GA to build the atlas. A total number of 89 scans of unique normal fetuses were included, with five to eight subjects in each week.

### dMRI data processing

The processing of dMRI data was performed in MRtrix3 and SVRTK. Raw data were preprocessed using a pipeline in MRtrix3 (https://www.mrtrix.org/), allowing for denoising, bias removal and between-volume motion correction.^36^ Images were then corrected for inter-slice motion and reconstructed to 1.2 mm isotropic resolution using SVRTK (https://github.com/SVRTK/SVRTK).^37^ This toolkit first performs iterative slice-to-volume registration and super-resolution reconstruction utilizing all 2D image slices regardless of gradient directions to generate a motion-free mean diffusion weighted (DW) 3D volume.^38^ A spherical harmonics model was then fitted to reconstruct all DW volumes with the corrected gradient tables.

We estimated diffusion tensors and obtained DTI-derived metrics including FA, MD, axial diffusivity (AD), and radial diffusivity (RD) using MRtrix3. FOD functions were calculated using CSD.^11^ Fixel-based analysis (FBA) were performed based on FOD to obtained fiber density (FD) and fiber-bundle cross-section (FC) for individual fiber populations within each voxel (fixels), which represent the microscopic axonal density per volume and morphologic fiber volume change with respect to the template, respectively.^39^ Fiber density and cross-section (FDC) was calculated by multiplying FD and FC to quantify the overall fiber content in WM regions.

### Atlas generation

We designed an FOD-based registration pipeline to reconstruct a population averaged template of the fetal brain for each gestational week (Figure 2). The pipeline consistent of two major steps of initialization and iterative registration.

**Figure 2.**
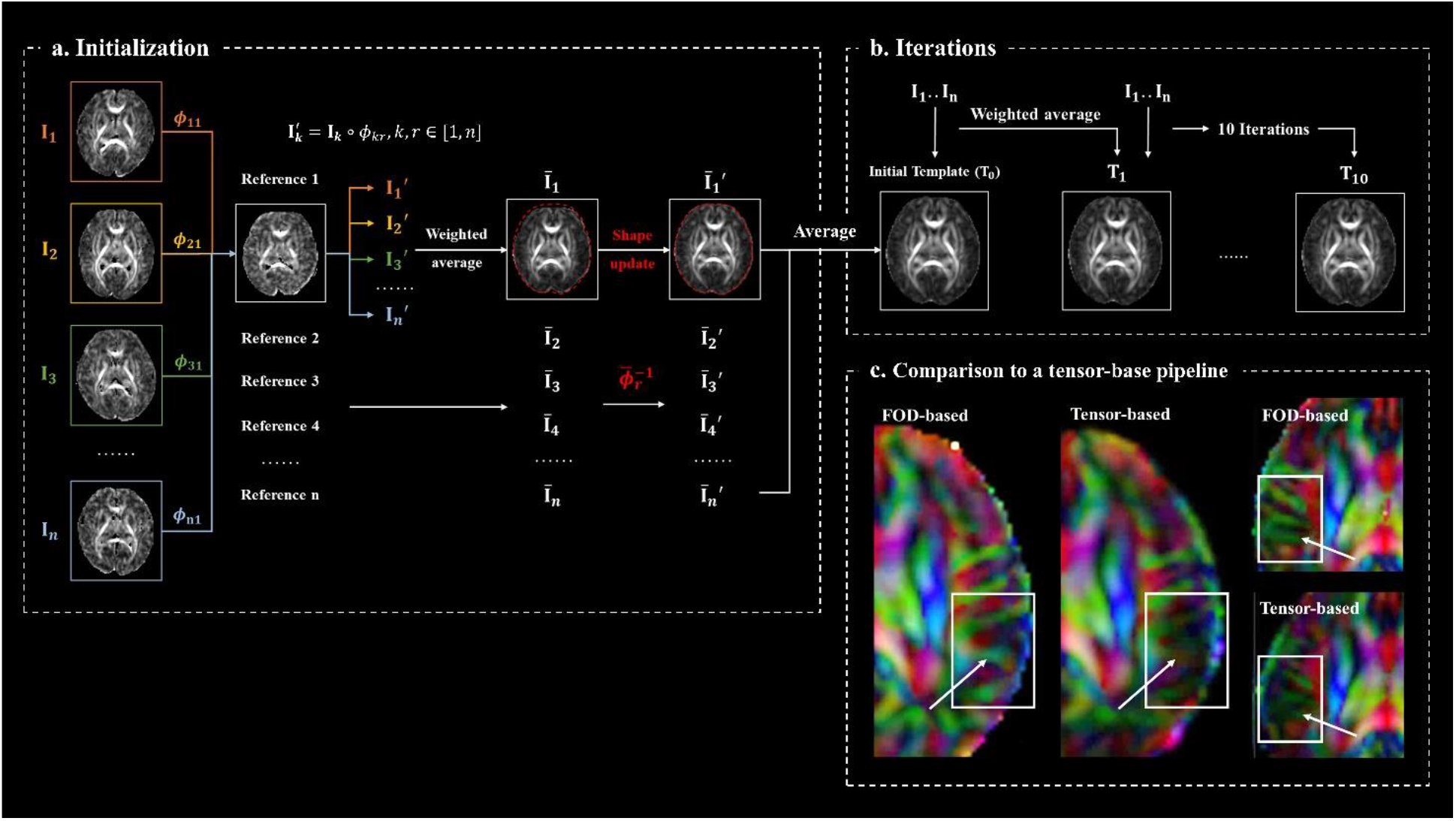
An FOD-based registration pipeline for generating fetal brain dMRI atlas. **(a)** For each GA, the template was initialized by pair-wise diffeomorphic registrations between subject FODs ***I_k_***(*k* ∈ [1, *n*]). The averaged images 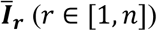 were obtained by calculating the weighted averages of the registered images 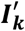, in which the weights were calculated according to GA. Shape updates were performed on 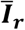 to get the unbiased 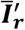. The initial template ***T*_0_** was then obtained by averaging the shape-updated images. **(b)** Subject FODs ***I_k_*** were then aligned to the template with FOD-based non-linear registrations, iteratively for 10 times and weighted averages of the co-registered subject images were calculated to obtain the final atlas. **(c)** The resulting atlas provides more structural details compared to the atlas constructed with tensor-based registration.

#### Initialization step

Proper initialization was critical to generate an unbiased population representation of the individuals, especially when the sample size is small. Here, the template was initialized by pair-wise diffeomorphic registrations between subject FODs.^40^ Iteratively for each subject image (***I_r_**, r* ∈ [1, *n*]), we took it as the reference and non-linearly registered the other images ***I_k_*** (*k* ∈ [1, *n*]) in the same GA group to the reference. GA-weighted average of the registered subject images 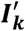 was computed to obtain the averaged images 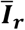 for each subject, according to Equations 1 and 2, where ***k*** was the index of the subject to be registered, ***r*** was the index of registration reference, ***n*** was the number of subjects in the GA group. The weight of each subject was calculated in each GA group using a kernel regression approach according to Equation 3, where *t* was the integer GA for each atlas (23W, 24W, etc.), and *t_k_* was the GA of subject ***k***.^41^

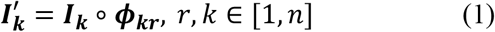

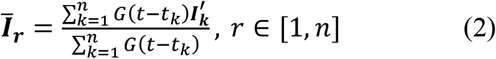

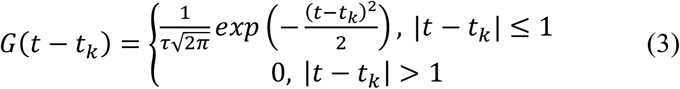

The initial template 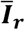 was biased towards the reference image ***I_r_*** as shown in Figure 2A, where the shape of 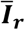 was biased from the averaged shape as indicated by the red dotted boundary. We eliminated the bias by applying the inverse of the averaged transform to the template.^28,42^ The generated unbiased template was 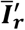:

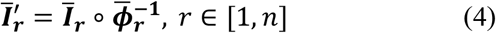

where

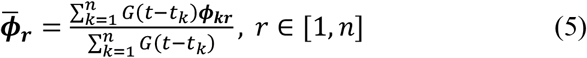

The initial template ***T*_0_** was then obtained by averaging the shape-updated images.

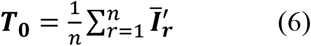

#### Iterative registration

Subject FODs were then registered to the initial template and averaged to obtain an updated template ***T*_1_**, and this process was repeated 10 times to obtain the final FOD atlas. The subject averages in each iteration were modulated by the kernel-regressed weights. All registration steps were performed in MRtrix3 with FOD reorientations.^43^ The final diffeomorphic transformations mapping from each subject to the atlas were obtained for generation of DTI atlas.

The DTI atlases were obtained by applying the nonlinear subject-to-atlas transformation from the aforementioned pipeline to diffusion tensors. The diffusion tensors of individual subjects need to reoriented before averaging, and this was performed using the *finite strain* algorithm:

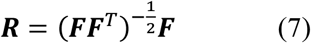

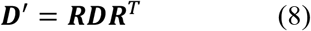

For each voxel, the rotation matrix ***R*** was calculated directly from the local Jacobian matrix ***F*** (Equation 7) and was applied to the transformed diffusion tensor ***D*** to obtain the transformed and reoriented tensor ***D***′ (Equation 8).^44^

### Fiber tracking

Probabilistic tractography were performed in the atlas space using the probabilistic algorithm iFOD2 in MRtrix3.^45^ Eight major WM tracts were estimated, including the genu, body and splenium of corpus callosum (GCC, BCC, SCC), cortico-spinal tract (CST), cingulum bundle (Cg), inferior longitudinal fasciculus (ILF), inferior fronto-occipital fasciculus (IFOF) and middle cerebellar peduncle (MCP). We registered MD maps of the atlas to the CRL fetal brain T2w atlas of corresponding GAs using nonlinear transforms, and back-transformed the CRL T2 segmentations to our atlas space for identifying regions of interest (ROIs). For each tract, one seed region was manually delineated on the colored FA map in atlas space. At least one manually drawn ROIs or T2 segmentation based regions were defined as inclusion or exclusion area. The delineation and selection of ROIs used in fiber tracking are described in the Supplementary Material. For unilateral tracts including CST, Cg, ILF and IFOF, streamlines passing the mid sagittal plane of the brain were removed. Other parameters for fiber tracking include: The threshold of FOD amplitude was set to 0.08, which is a bit higher than the commonly used threshold value in adults (default = 0.05), in order to reduce the false positives; the angular threshold was set to 15° for CST, IFOF, and ILF, and 20° for CC, Cg, and MCP that have larger curvatures. After obtaining the original fiber tracts through automatic probabilistic tracking, we used the Topographic Tract Filtering tool to remove spurious streamlines that deviated from the trunk of tract.^46^ We used the *group graph spectral distance* algorithm in this toolbox and the parameters were adjusted for different tracts to achieve the desired quality. The tracts were transformed back to subject spaces using the inversed subject-to-atlas transformations resulted from the atlas generation pipeline.

### Statistical analysis

The mean FA, MD, AD, RD, FD, FC values along the WM tracts were extracted from each subject. To map fixel-based values (FD and FC) to voxel space, we took the sum of fixels in each voxel. For CST, IFOF, ILF and Cg, the values were calculated by averaging the measurements from bilateral areas. We further segmented CST into four segment from inferior to superior regions with equal length to examine the developmental patterns along the tract. DTI- and FBA-based measurements of WM tracts were extracted from individual subjects to fit a flexible sigmoid function over GA in MATLAB (The MathWorks, Inc.), except for FC, which was linearly fitted.^47^ The AD and RD values were Z-scored before sigmoid fitting so that they can be directly compared. The flexible sigmoid function is given in Equation 9:

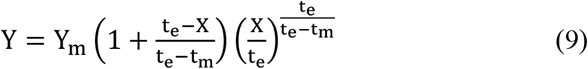

The model gives a turning time point t_e_ for non-monotonic data, which in our case marked the time when each value reached their peak or valley. Y_m_ is the peak value of the fitted curve and t_m_ is the time of a mid-way inflection point, when the curve has the highest growth rate.

Fixel-wise regression analysis of FD, FC, FDC with GA was performed to evaluate the GA-dependencies of these measurements. The analysis was performed within the FBA pipeline using MRtrix3. First, FC and FDC fixel maps of all subjects from 24 to 38 weeks were registered to their population-averaged template in fixel space, to enable fixel-wise statistics in a common space. Linear regressions were performed between GA and FD/FC/FDC, in a fixelwise manner. For FD, we divided the subjects into two groups with fetuses before and after 31W, to study these two periods separately, as the different maturation patterns of WM microstructures can lead to non-linear trends of FD before and after 31 weeks of gestation.^6,26^ The whole-brain statistics underwent 5000 permutations with the connectivity-based fixel enhancement, which is a threshold-free cluster enhancement-like approach using fiber connectivity information as a prior to smooth and enhance the statistic maps.^48,49^

### Group analysis with CHD subjects

Comparison between normal and CHD fetal brains was performed with ROI-based analysis. We focused on GCC and SCC of the fetal brain, as previous studies have reported volumetric and microstructural abnormalities in CC of the fetuses and children with CHD.^50,51^ ROIs of GCC and SCC were manually defined on the 31W colored FA map (Fig. 8a) and non-linearly transformed to other GA spaces. We referred to the WM segmentations in JHU single subject neonatal atlas for the manual ROI delineation.^52^ We extracted the ROI-averaged DTI measures from CHD and normal fetuses. Analysis of Covariance (ANCOVA) was performed between CHD subjects (n = 11, GA 25-30 weeks) and GA-matched normal subjects (n = 40, GA from 25 to 30 weeks) with GA as a covariate in MATLAB using the GRETNA 2.0 toolbox (https://www.nitrc.org/projects/gretna/). We also compared the 28W subjects separately, including eight normal subjects and five CHD patients.

## Results

### The spatiotemporal fetal brain dMRI atlas

A spatiotemporal dMRI atlas of the Chinese fetal brain at GA from 24 to 38 weeks were generated using the proposed pipeline, containing both the DTI- and FOD-based atlases (Figure 3). Compared to the existing fetal brain dMRI atlas generation pipeline with tensor-based registration, the atlas generated by the proposed pipeline demonstrated sharper structural details in cortical regions (arrows in Figure 2C).

**Figure 3.**
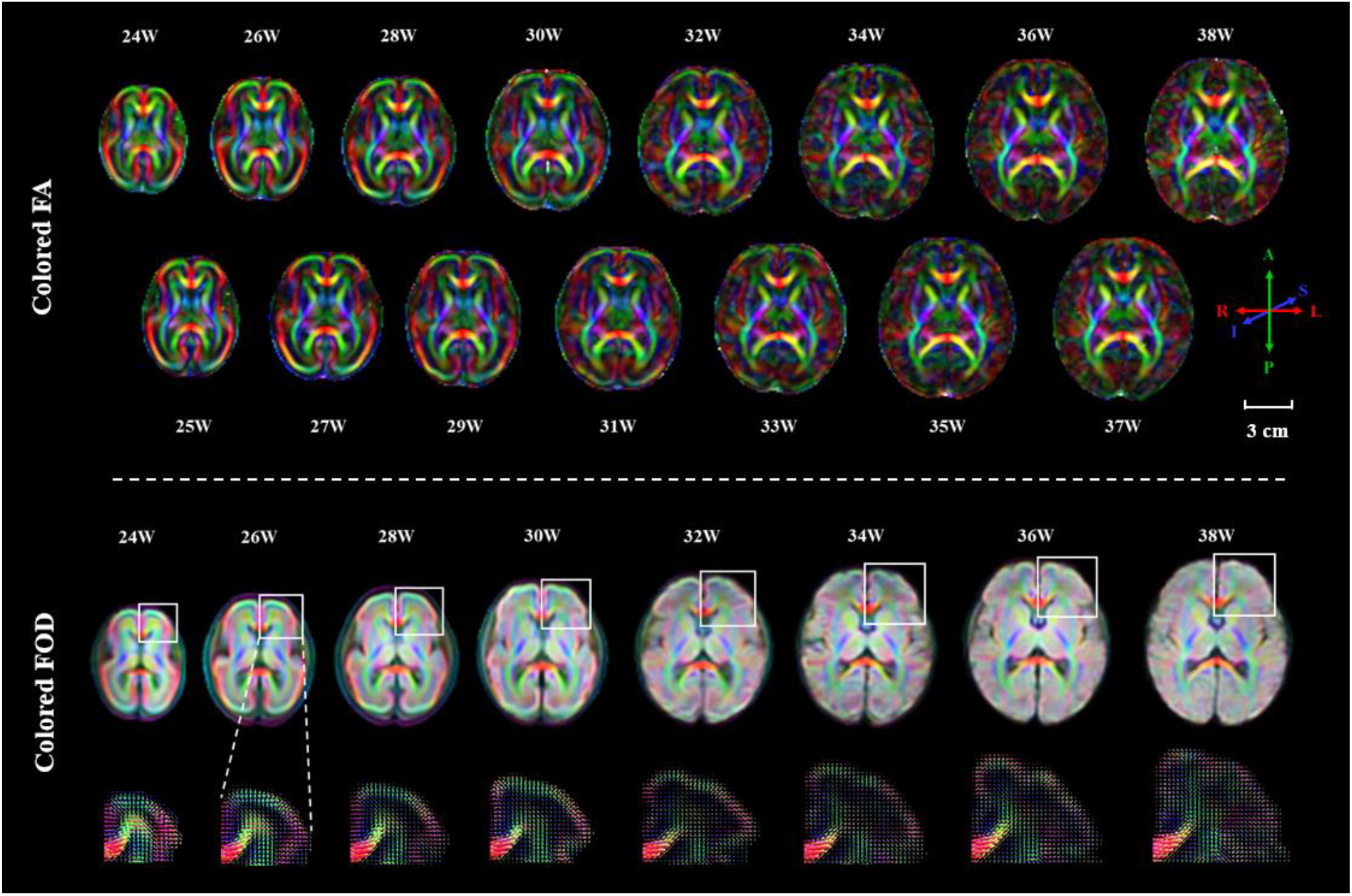
Colored FA and colored FOD templates from the fetal brain dMRI atlas of gestational age from 24 to 38 weeks. The atlas revealed that the fetal brain experienced significant microstructural development during gestation. The zoomed-in FOD profiles showed the loss of radial coherence in the cortical plate (CP) over GA as the cortex develops into more complex structures.

The atlas revealed the dramatic structural alterations of the fetal brain from 24 to 38 weeks of gestation. In the colored FA maps, only a few major WM bundles are discernable in the younger fetal brains, e.g. at 24-26 weeks. More detailed and complex fibers structures can be observed in later GAs. The zoomed-in FOD profiles in Figure 3 reflected the unique developmental patterns of cortical regions. The cortical microstructural were well-organized in the radial direction at 24 weeks and gradually became multi-directional with more fiber crossings, indicating the loss of radial coherence in the cortical plate over GA as the cortex develops into more complex structures. The population atlas is deposited for public use (https://github.com/RuikeChen/Fetal-Brain-dMRI-Atlas).

### Developmental pattern of major WM tracts

Nine WM tracts were successfully estimated in the atlas space from 24 weeks to 38 weeks, as displayed in the sagittal view at representative GAs of 24W, 27W, 30W, 33W, and 36W (Figure 4), which exhibited unique morphological development. For instance, the projection fibers, commissural fibers, and brainstem fibers were relatively well developed starting from 24 weeks, while the associated fibers, such as IFOF and SLF, and limbic fibers of Cg gradually grew over time with more widespread extensions at later gestation. The SLF seemed to be the latest to develop that cannot be tracked until 25 weeks, similar to Huang, et al. ^53^ from *ex vivo* samples.

**Figure 4.**
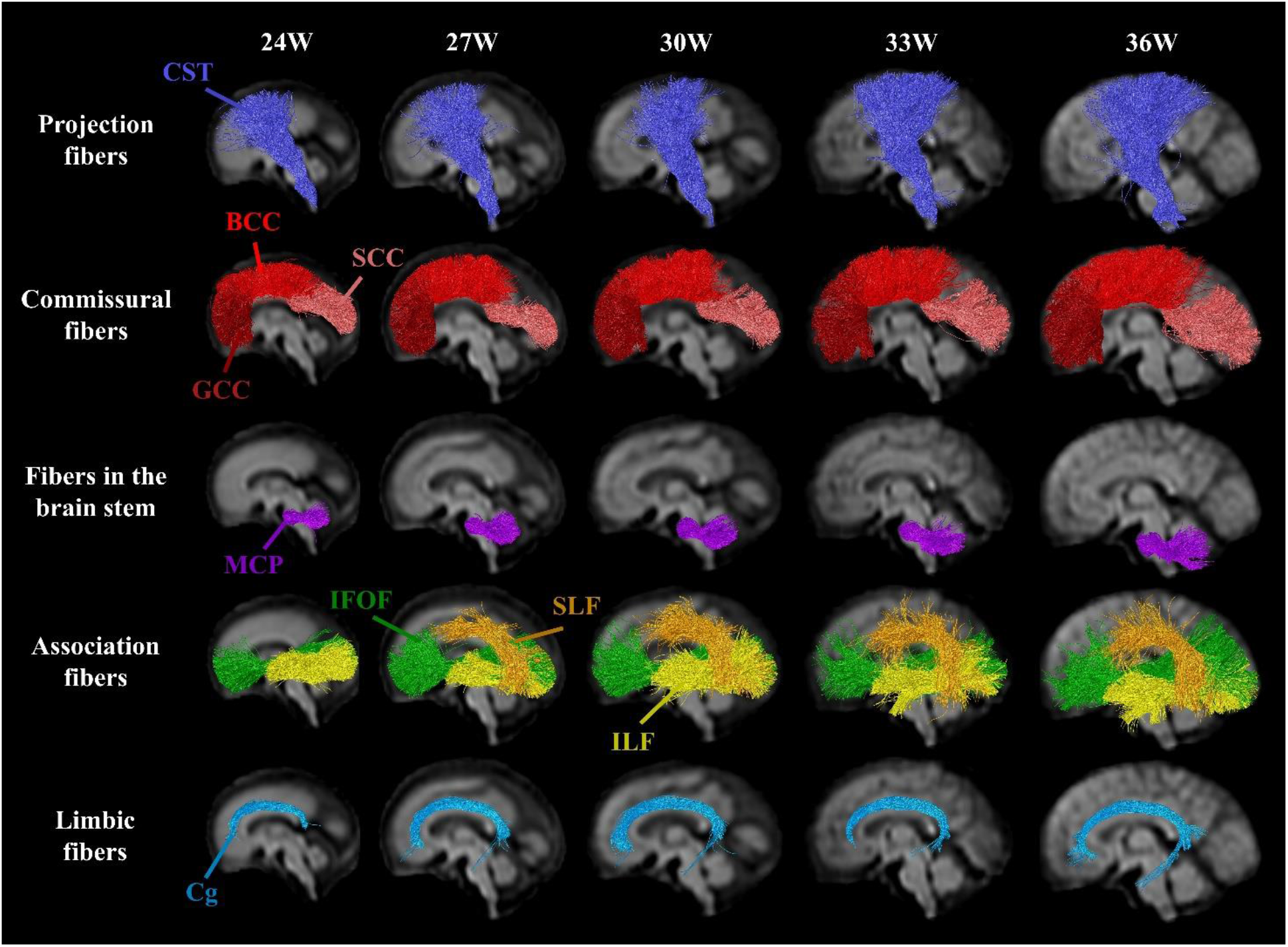
White matter fiber tracts reconstructed from the fetal brain dMRI atlas. WM tracts including Cortico-spinal tract (CST), genu, body and splenium of corpus callosum (GCC, BCC, SCC), middle cerebellar peduncle (MCP), inferior fronto-occipital fasciculus (IFOF), superior longitudinal fasciculus (SLF), inferior longitudinal fasciculus (ILF), and cingulum bundle (Cg) are displayed in representative GAs, showing unique morphological development. All tracts were successfully estimated except SLF, which could only be traced after 25 weeks of gestation.

The tract-averaged developmental trajectories of dMRI measurements were depicted in Figure 5a. We found the MD values in most WM tracts increased first with slightly decreasing FA values; and after around 30W, the tracts exhibited a decrease of MD and increase of FA. The turning points of FA and MD in the eight WM tracts (excluding SLF) given by the flexible sigmoid growth function were marked in Figure 5a, which were statistically significant in almost all the curves (*P* < 0.05), except for the FA trajectories of Cg (*P* = 0.7218) and SCC (*P* = 0.1733). The tract-specific developmental trajectories of FA and MD can be summarized as below:

1. MD and FA of MCP changed monotonically from 24W to 38W, indicating that the turning points were probably before 24W.
2. CST had an early turning point in both MD (26.46W) and FA curves. The MD values in the four segments of CST also displayed different patterns (Figure 6b). Segment 1 and 2, which are the inferior parts close to the brain stem, showed continuously decreasing MD from 24 to 38 weeks, indicating the turning points were earlier than 24 weeks. Segment 3 and 4, corresponding to the superior corona radiata and the posterior limb of internal capsule, reached their peaks MD values at around 26 and 28 weeks, respectively. Therefore, the turning points of MD along the CST demonstrated an inferior-to-superior order.
3. MD of the cingulum bundle reached a peak at 27.09W but FA of this bundle did not show a statistically significant turning point.
4. The association fibers including the IFOF and ILF had late turning points of MD at 29.23W and 29.52W. The turning points of their FA values were at 26.54W and 30.31W respectively.
5. The turning points of MD and FA in CC varied in its different segments. The turning-point of MD occurred first in SCC (27.47W), followed by the body (30.57W), and then the genu (33.45W). FA in SCC increased from 24W to 38W, while BCC and GCC had turning points of FA at 29.55W and 32.01W, respectively. The change of both MD and FA in CC showed a posterior-to-anterior order.

**Figure 5.**
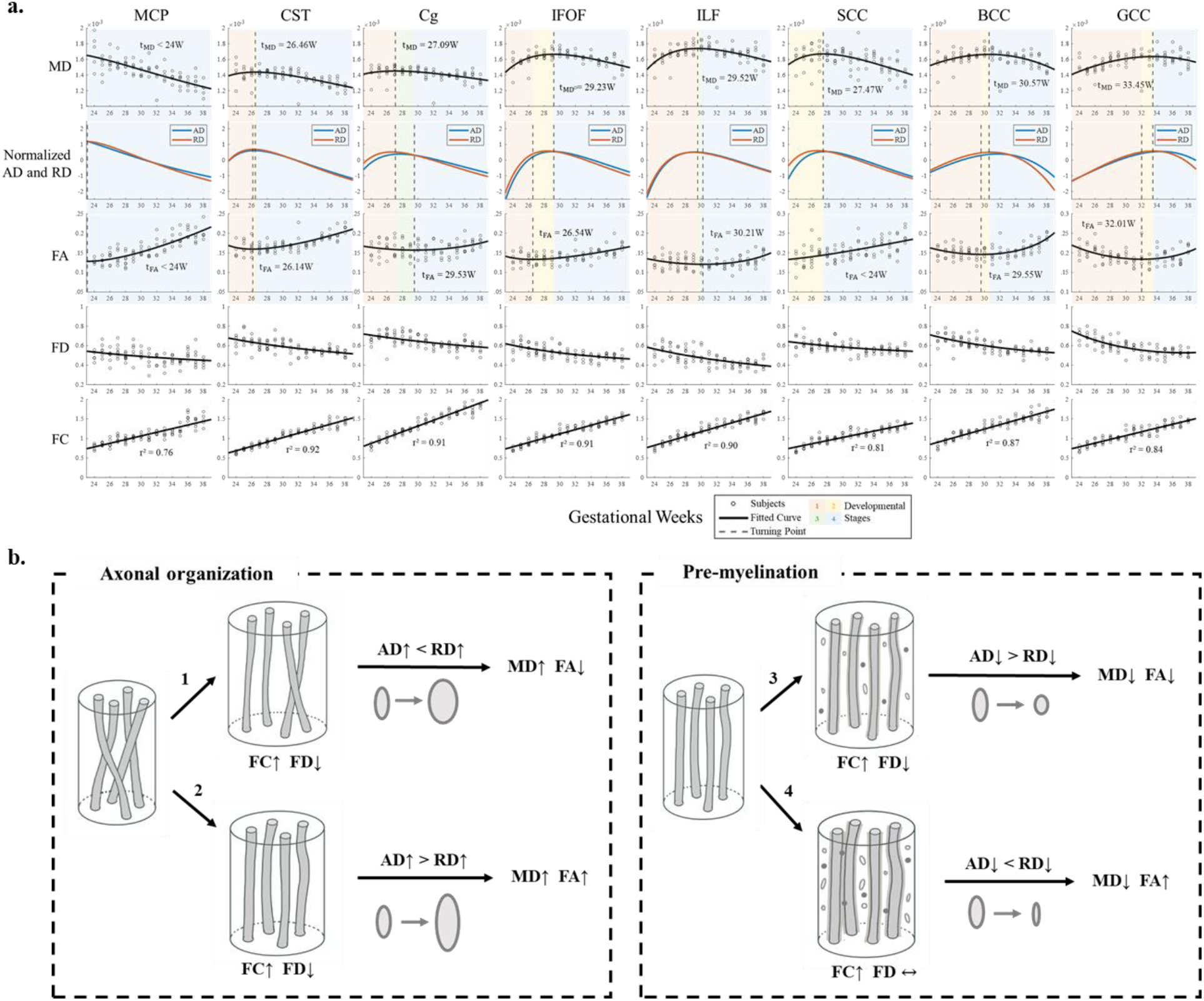
The developmental trajectories of major WM tracts. **(a)** Trajectories of dMRI measurements in eight major WM tracts. **(b)** Based on the relative changes of the multi-modal dMRI metrics, we hypothesized four phases with distinct microstructural events: 1) fast volumetric growth with relatively slow axonal organization, characterized by a more rapid increase of RD than AD, leading to the initial increase of MD and decrease of FA; 2) fast fasciculation of axons that outgrows the effect of volume expansion, indicated by the faster increase of AD than RD, resulting in simultaneous increase of MD and FA; 3) Slow pre-myelination and growing of WM volume at the same time, corresponding to increasing FC and relatively unchanged FD. AD decreases more than RD; 4) Fast pre-myelination when immature oligodendrocytes proliferate rapidly and contact with axons to form pre-myelin sheaths, reflected by increasing FD and FC. RD decreases more than AD during this phase and MD decreases while FA increases. The four developmental phases are represented by different colors in **(a)**.

**Figure 6.**
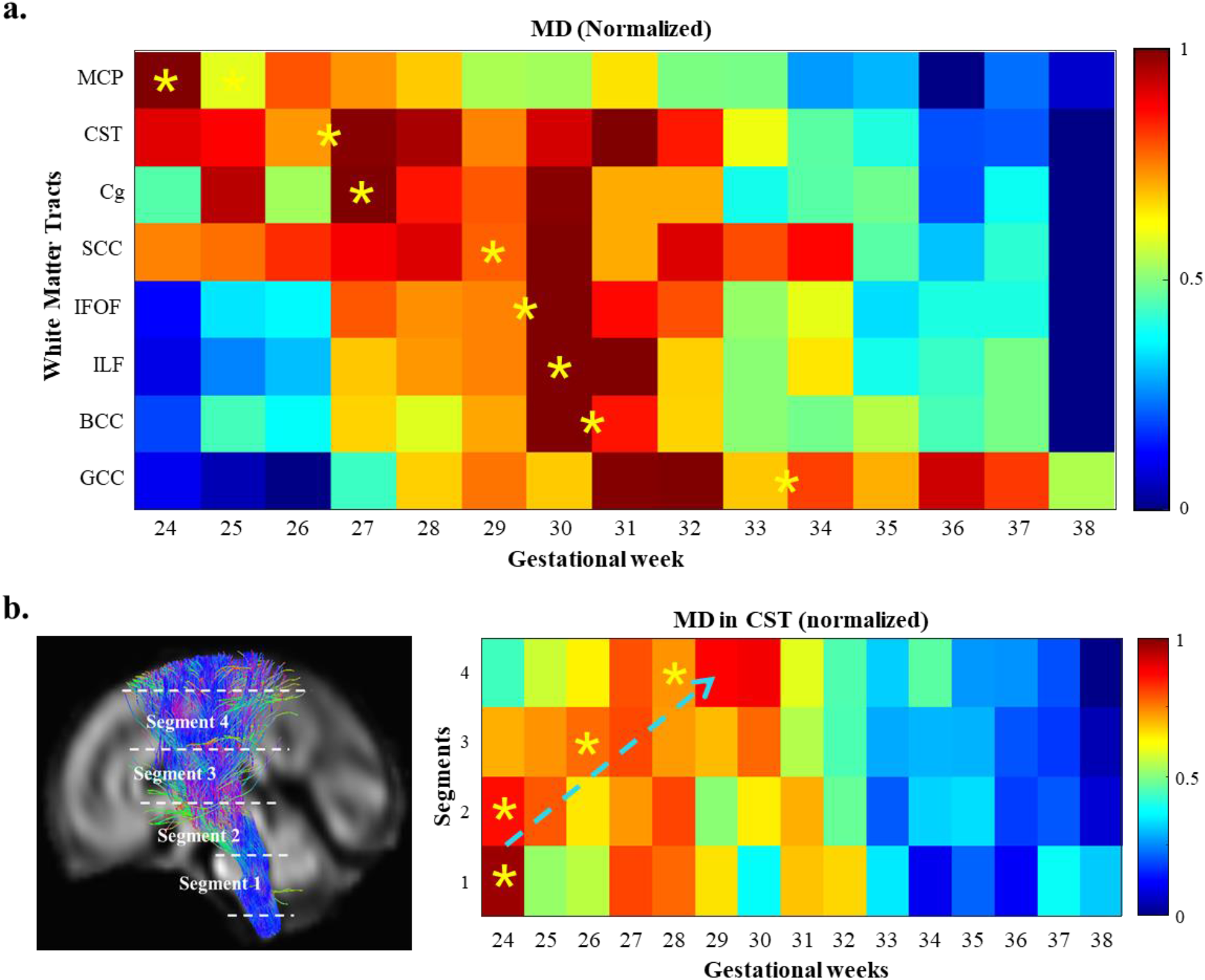
The developmental order of WM tracts revealed by the turning points of the MD trajectories. (**a)** The heat map displays the normalized MD values of the eight target WM fiber bundles, where the turning points given by the flexible sigmoid growth function-fitting are asterisked. (**b)** The CST was divided into four segments with equal length from inferior to superior positions. The turning-points of the MD curves in each segment were indicated in the heatmap, following an inferior-to-superior order.

Both AD and RD had similar trends as MD in these eight WM tracts but their relative changes varied during different stages. FD decreased at first but the rate slowed down at around 30W, and was almost stable afterwards. FC in all studied tracts significantly increased over GA with *R*^2^ ≥ 0.75, indicating linearly increasing fiber volume (Figure 5a).

Given the non-linear trajectories of these measurements and the fitted turning points of MD and FA, we hypothesized that these changes were driven by distinct developmental events at different stages. We divided the trajectories into four phases according to the turning points of MD and FA, which were indicated by different colors in Figure 5a. Phase 1 is characterized by increasing FC and decreasing FD, leading to the initial increase of MD and decrease of FA, which together indicate fast WM volumetric growth with relatively slow axonal organization. In phase 2, MD and FA increase simultaneously, indicating the fast fasciculation of axons outgrows the effect of volume expansion. During phase 3, FC continues to increase and FD shows a slower decrease. Both MD and FA decrease in this phase, which suggest the slow pre-myelination and fast growing of WM volume occur at the same time. Phase 4 is identified by increasing FC and relatively unchanged FD. MD decreases while FA increases during this phase, indicating the immature oligodendrocytes begin to proliferate rapidly and contact with axons to form pre-myelin sheaths. Phase 1 can be observed in most of the fiber tracts, except for MCP and SCC, and we assumed that phase 1 has ended by the time of 24W for these tracts. The intermediate phase 2 and phase 3 selectively occur in different tracts, depending on the relative rates of fiber volume increase and axonal fasciculation/pre-myelination, indicated by the colored phases overlaid on the developmental trajectories. Phase 4 can be observed in all studied tracts as it begins in the late second or the third trimester and continuous toward 38 weeks of gestation.

According to this hypothesis, the turning points of MD mark the beginning of pre-myelination. Mean MD values of each WM tracts over GA were normalized and displayed together in a heat map with (Figure 6a), and the order of turning points agreed with the developmental order of WM myelination reported in previous studies.^4,7,54^ The developmental order along the different segments of a fiber tract can be also quantified in a similar manner, e.g., an inferior-to-superior developmental order was found along the CST (Figure 6b).

### FBA of GA-dependent microstructural changes

FBA-based statistical analysis reveals significant microstructural changes in a voxelwise manner. Figure 7a-b showed bran regions with significant increase of FC and FDC over GA. The intensities were scaled by the growth rate of corresponding values. It is observed that the center of WM tracts exhibited the fastest growth, and FC also showed volumetric expansion in the cortical regions. FD, however, significantly decreased before 31 weeks of gestation in GCC, CST, and ILF, as displayed in Figure 7c, possibly due to the fast growth of FC that outperformed the effect of axonal fasciculation or pre-myelination. After 31 weeks, no significant change of FD was found. In addition, we found CC displayed unique developmental patterns in its genu, body and splenium parts, where SCC had the most significant changes of FC, FD and FDC (Figure 7d).

**Figure 7.**
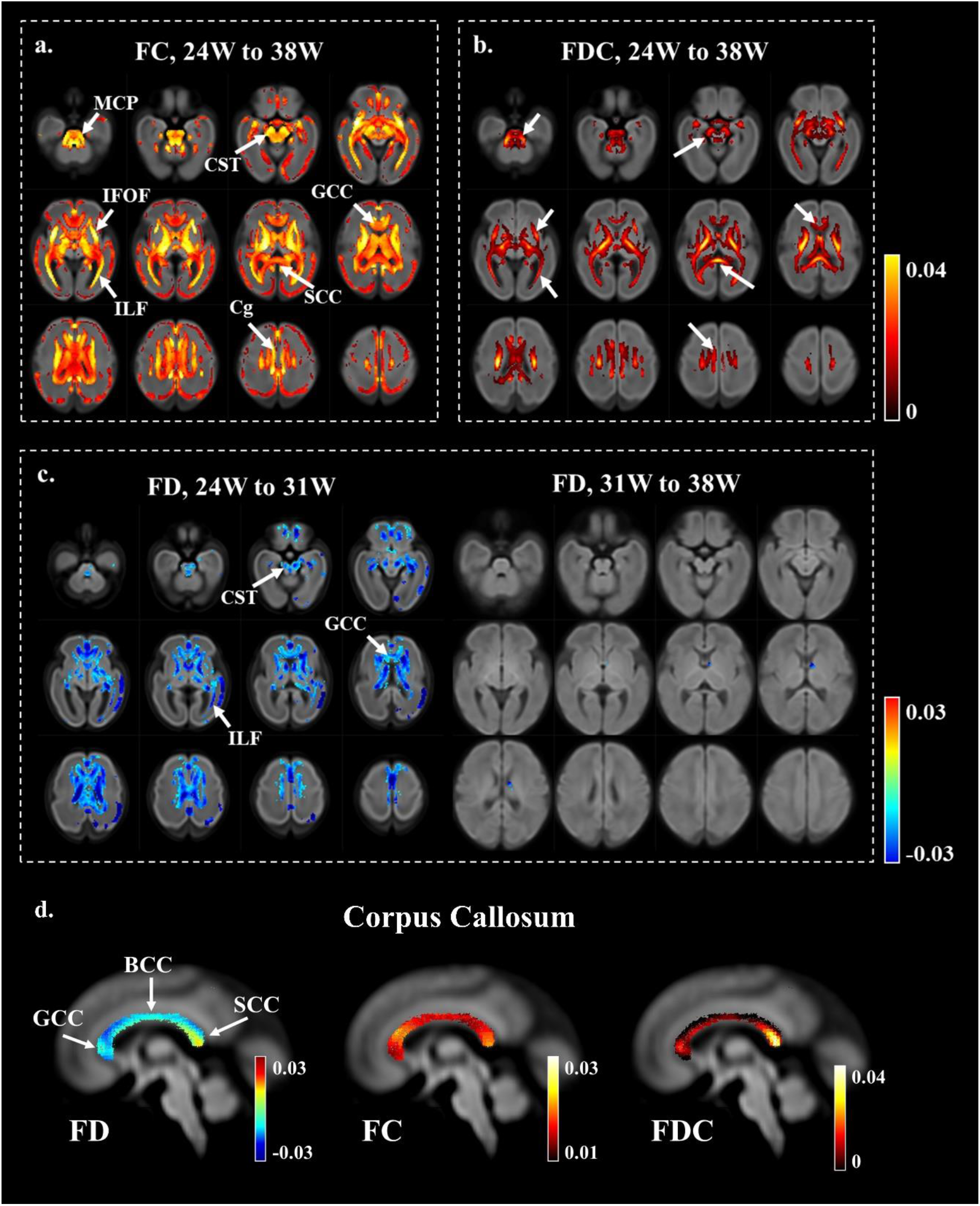
Fixel-based analysis (FBA) of the GA-dependent microstructural changes in a voxelwise manner. Regions with statistically significant change (corrected *P* < 0.05) are overlaid on the FOD image and the colors indicated the growth rate in each voxel. **(a-b)** FC and FDC values increased significantly in MCP, GCC, SCC, ILF, IFOF, CST, and Cg during 24W to 38W. **(c)** FD significantly decreased before 31 weeks of gestation in GCC, CST, and ILF, and had no significant change after 31W. **(d)** Voxelwise growth rate in FD, FC, FDC in the corpus callosum. SCC had the highest growth rate among all three measurements.

### Altered WM microstructures in the brains of fetuses with CHD

Group comparison revealed that CHD fetuses had significantly lower FA values in SCC (*t* = −2.4134, *P* = 0.0197) and GCC (*t* = −2.1993, *P* = 0.0327) compared to normal subjects between GA of 25-30 weeks. Figure 8a displayed the group averaged FA after regressing out the GA effects as well as the raw FA values for each individual subjects. We also examined 28W-GA subjects separately, and found CHD fetuses had significantly higher MD values in SCC (*t* = 2.2560, *P* = 0.0454) and no significant group difference in GCC in the 28W-GA fetus.

**Figure 8.**
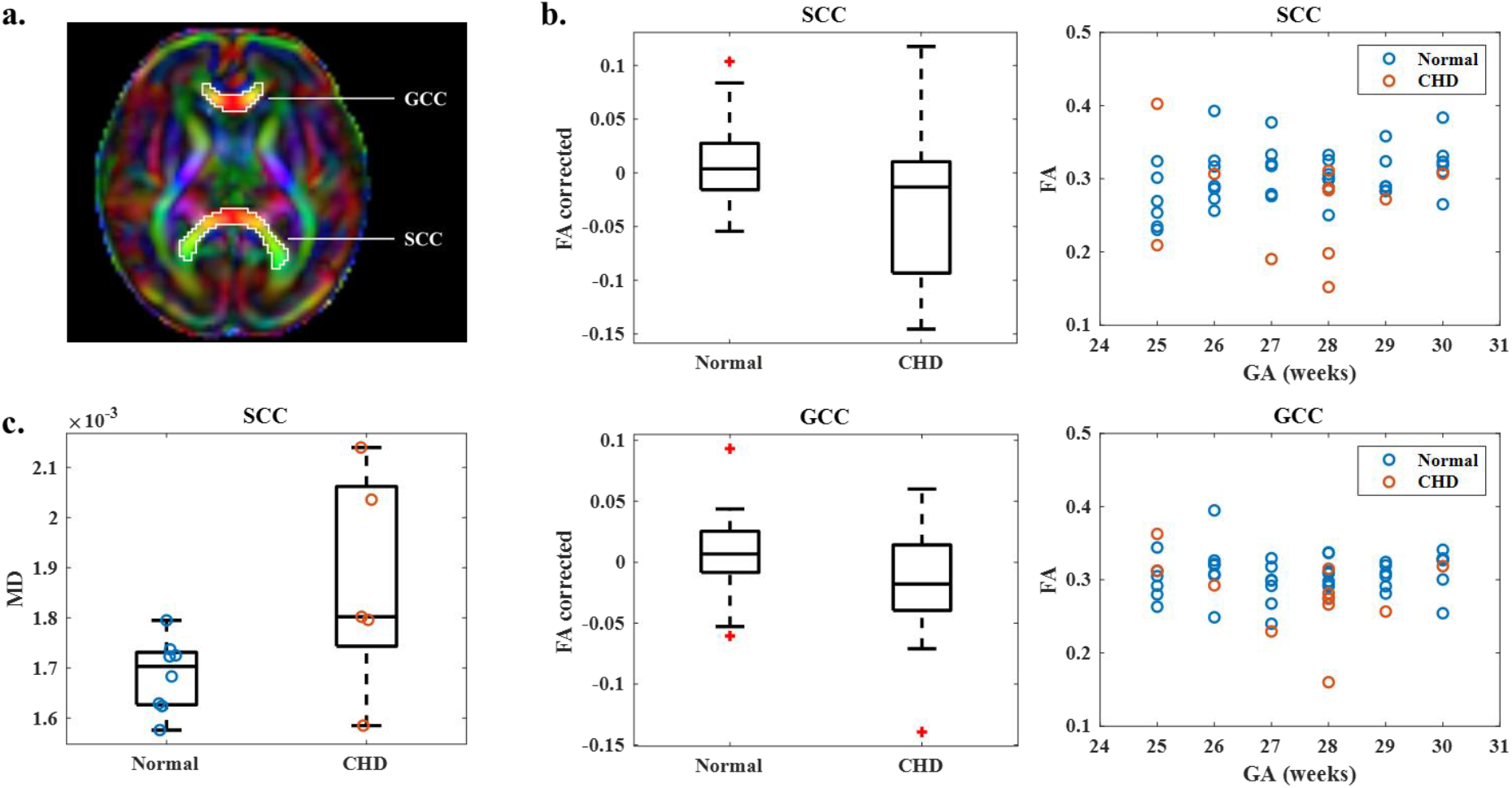
Group comparison with ROI-based analysis between normal and CHD fetal brains. **(a)** ROI definition of GCC and SCC on the 31W colored FA. **(b)** Analysis of Covariance (ANCOVA) with GA as a covariate was performed between CHD fetuses (n = 11) and GA-matched normal subjects (n = 40, GA = 25W ~ 30W). The results showed significantly reduced FA in SCC and GCC of the CHD fetal brains compared to normal fetuses (*P* < 0.05). Both the group averaged FA values after regressing out the GA effects and the raw FA values of individual subjects were shown. **(c)** The 28W subjects were compared separately, including eight normal subjects and five subjects with CHD. Compared to normal fetuses, the diseased fetal brains showed significantly increased MD in SCC.

## Discussion

In this study, we generated a spatiotemporal dMRI atlas of the Chinese fetal brains from 24 to 38 weeks of GA for the first time. We achieved better registration accuracy than the existing fetal brain DTI atlas by using used an FOD-based pipeline initialized with pair-wise registrations. The atlas includes both DTI-based and FOD-based metrics to provide higher-ordered diffusion information of the fetal brain than the existing DTI atlas, which revealed non-monotonic changes of dMRI measurements over GA in the major WM tracts.^28^ Compared to previous studies that mainly utilized DTI-based parameters to depict the WM development of the fetal brain, we jointly analyzed the DTI- and FBA-based measurements that gave rise to a hypothesis of distinct developmental phases.^26,27^ We divided the non-linear trajectories of dMRI measurements into four phases according to the characteristic changes of MD/FA/FD/FC and corresponded them to the known microstructural events.^25,55^ This hypothesis explained the dMRI patterns more comprehensively by considering both volumetric and microstructural growth, and specifically pointed out the turning point of MD may indicate the start of pre-myelination. The developmental order derived from this hypothesis agreed with the order of myelination in histological findings.^4,56^

### Spatiotemporal characteristics of the fetal brain dMRI atlas

In this work, we generated the first dMRI atlas of the fetal brain in a Chinese population using a FOD-based atlas generation pipeline. As visualized in Figure 2, the atlas generated by the proposed FOD-based pipeline demonstrated more structural details in cortical regions compared to the existing tensor-based pipeline, indicating the higher inter-subject registration accuracy with the high-order FOD information.^28^ It is known that cortico-cortical association fibers emerge in the fetal brain starting from 30 weeks of gestation; since then, the initial radial organization of the cerebral mantle became less prominent due to the development of complex cortical connectivity.^57,58^ These patterns were captured in the zoomed in FOD profiles in Figure 3, where the radial coherence in the cortical plate gradually disappeared and crossing fibers emerged over GA. Therefore, the diffusion tensor model may not be sufficient in representing the complex microstructures in the fetal brain. The use of high-order dMRI does not only model the cross-fiber to improve the registration accuracy, but only provide additional microstructural contrasts in the dMRI atlas that was jointly used in this study to staging the WM development. Furthermore, we utilized pair-wise registrations with shape updates in the initialization of the atlas generation, which is important to avoid bias from individual brains, especially when the sample size is small (*n* = 5-8).

Prior to this study, the only reported fetal brain DTI atlas was the CRL atlas, which was generated from a Caucasian fetal population. The morphological difference between the Chinese and the Caucasian brain can be observed as early as six year old.^32^ It would be ideal to directly compare the Chinese fetal dMRI atlas with CRL atlas, which however, is difficult due to the different data acquisition, reconstruction, and atlas generation pipelines between two atlases. For the CRL atlas, subject data were constructed from multiple stacks including the axial, coronal and sagittal views to enhance the through-plane resolution. However, it has poor in-plane spatial resolution compared to our atlas (Supplementary Fig. 5), possibly because of the lower in-plane resolution (2 mm) and poor angular resolution (12 gradient directions) in data acquisition and the limitations in atlas generation as discussed above. A visual comparison of DTI-derived values between the two atlases can be found in the Supplementary Material. The developmental trends of FA and MD were similar between the two atlas but the absolute values differed (Supplementary Fig. 6). The difference may both come from the biological and technical factors. We further investigated the effect of super-resolution reconstruction on fetal brain dMRI data and found the reconstruction using single 2D stacks and multiple 2D stacks gave slight different FA and MD (Supplementary Fig. 7), which may partially explain the atlas difference we observed. Therefore, to study the biological difference between the Chinese and Caucasian during fetal stage, more strictly designed experiments are needed.

We also demonstrated the potential utility of the normative atlas in diagnosis of CHD, and found significant different measurements in GCC and SCC. The significant lower FA and higher MD values in these WM tracts may reflect the delayed brain maturation in CHD fetuses as reported before.^59,60^ Particularly, the early myelinated WM tracts with high metabolic demands may be more vulnerable to insufficient supply of substrates in CHD fetuses.^61^

### Early development of WM tracts

Eight WM tracts including the major projection, commissural, association, and limbic fibers were successfully estimated from 24W to 38W fetal brain atlas except SLF, which could only be traced after 25W. Complex nonlinear trends of dMRI measurements were observed in these WM tracts, with an initial decrease of FA and increase of MD before around 30W, followed by opposite trends. The results pointed to an important transition in developmental stages at around 30W of gestation, when active neurological events such as pre-myelination takes place.^1^ The non-linear changes of MD and FA were also reported in recent studies.^26,27^ Wilson, et al. ^26^ studied 113 fetuses at GA from 22 to 37 weeks and found initial decrease of FA and increase of MD before around 30W, followed by reversed changes, and associated the increase of FA with pre-myelination. But the relationship between the trajectories of MD and FA was not explained and microstructural features beyond the tensor model were not considered; and the initial decrease of FA was not fully understood. Similar results were found later in the CRL atlas using a deterministic tractography method, and the authors related the initial decrease of FA to the growth and involution of the transient compartments in the fetal brain.^27^ The current study offered a different interpretation of this trend by joint analysis of the microscopic and morphological changes of WM.

Compared to previous studies, we quantified the non-linear trends to a flexible sigmoid growth function to identify critical turning points. Note in our results that FA and MD did not always change in an opposite manner, that is, the turning times of FA and MD can be different in a same WM tract. The turning points also varied between different WM structures. These patterns suggested that the WM development of the fetal brain underwent several biological processes with a critical turning point in the second trimester around 30W. During the second trimester, the WM fiber bundles of the fetal brain are still quite underdeveloped and are rich of extracellular matrix (ECM).^62^ Although oligodendrocytes (OLs) have already began to proliferate and differentiate, but the process is slow and active myelinogenesis does not start due to the “arrested” state of immature OLs, indicating a relatively slow microstructural development during this period.^6^ On the contrary, the volumetric growth is active during the second trimester. As displayed in Figure 7a, FC and FDC significantly increased over 24W to 38W of gestation (*P* < 0.05), indicating the volume and the total amount of axons in WM fibers grew rapidly. FD values, however, decreased before 31 weeks and had insignificant change afterwards. Increasing FC and decreasing FD suggests that the relative density of axons per volume declined due to the fast volume expansion and relative slow microstructural change. The inconsistency between microstructural and volumetric growth in WM tracts has been reported in a previous study that IFOF and ILF had rapid growth in volume but changed slowly in DTI measures.^21^ A recent study explained that the expansion of subplate contributes to the increase of ECM in WM tracts during the second trimester, which also suggested the volumetric growth can result in reduced microstructural complexity.^27^

### Biophysical interpretation of the developmental trajectories

The WM development of the fetal brain was separated to three stages based on both DTI and biological findings in, following the order of axonal organization, proliferation of OLs, and myelination, which was matched to the nonlinear changes in FA and MD.^25,55^ However, they did not observe the initial decrease of FA in WM bundles as reported in this and other recent studies, maybe due to the limited number of subjects involved.^26,27^ The theory ignored the change in volume and total amount of axons in WM bundles; and also, the third order polynomial function the authors adopted is not optimal for characterizing growth curves, or for dividing developmental stages. In the view of the multi-modal dMRI-based developmental pattern uncovered in the current study, we hypothesize four phases corresponding to distinct microstructural events, considering both volumetric and microstructural changes of the axonal bundles (Figure 5a). The starts and ends of these phases are marked by the turning points of MD or FA (Figure 5b).

#### Phase 1

Quick volumetric growth with relatively slow axonal organization. During this stage, the intertwined axons became better aligned in the axial direction, resulting in an increase of AD. On the other hand, the significant volume expansion in the radial direction resulted in a faster increase of RD than AD, which in turn resulted in an increase of MD and decrease of FA. This stage ends when either MD or FA reached its turning point.

#### Phase 2

On the other hand, it is possible that the axonal organization is fast that it contributed more to DTI measures than the volumetric growth. In this case, AD increases faster than RD, and MD and FA increase simultaneously, which was observed for SCC in the current study. This stage starts from the turning point of FA and ends at the turning point of MD.

#### Phase 3

The transition from the axonal organization stage to pre-myelination stage result in the decrease of diffusivities. The increase of immature OLs in WM tracts leads to the reduced ECM, which contributes to the decrease of both AD and RD. At the beginning of pre-myelination, however, microstructural maturation are slow due to the “arrested” stage of immature OLs, and the radial volume expansion remains the major factor contributing to the changes of DTI-derived parameters.^6^ In this situation, RD decreases less than AD, resulting in simultaneous decrease of MD and FA. This stage starts when MD starts decreasing and ends at the turning point of FA.

#### Phase 4

When the immature OLs start to proliferate rapidly and contact the axons to form pre-myelin sheaths, RD is more hindered by these sheaths and decreases more than AD. During this period, MD decreases while FA increases. This stages starts after the turning points of MD and FA, and continues till birth.

In phase 1-3, FC increases while FD decreases significantly, due to the fast volumetric growth and relatively slow microstructural maturation and the relative content of axons in each voxel decreases due to the quick volume expansion. In phase 4, FC continues to increase and FD value becomes stable due to the balanced changes of pre-myelination and volumetric growth. Phase 2 occurs for some of the WM tracts when the turning point of FA is ahead of MD, and phase 3 occurs in the opposite situation. According to this theory, the turning point of MD marks the beginning of pre-myelination in WM fibers, when the axons fasciculation is almost finished and the glial cells start to proliferate and differentiate. Initial ensheathment of axons by immature OLs starts later to prepare for true myelination.

By comparing the model-fitted turning points of MD in the eight major WM tracts, our finding suggested that pre-myelination starts earliest in the brainstem, followed by projection fibers and limbic fibers, and the commissural fibers and the association fibers showed late onsets. This order agreed with the previously reported order of WM myelination by previous histological studies.^4,7,54^ Major WM pathways such as the CST and CC exhibited within-tract variability with a proximal-to-distal and posterior-to-anterior sequence of pre-myelination, respectively. CC demonstrated with a wide range of turning-points from 27.47-33.45 weeks. As reported before, although GCC grows faster than SCC in early gestation, myelin structures were discovered in SCC earlier than GCC after term.^4^ Interestingly, we found the GCC had a turning point of FA close to MD, whereas FA value in SCC linearly increased from 24W to 38W. Moreover, FA in SCC was lower than GCC at the beginning but increased to a comparable level near term, and the pattern was also observed in previous dMRI studies, supported by the evidence that posterior part of CC is less developed than the anterior part at around 20W.^26,28,53^ According to our theory, the simultaneous increase of FA and MD indicated that SCC experienced quick axonal organization in phase 2, which also suggested the delayed but fast-paced development of SCC compared to GCC. In addition, we found Cg had an early turn of MD (27.09W) but the turning point of its FA value was not statistically significant. Although limbic fibers emerged early, as described by *ex-vivo* studies, myelin structures were seen after term in histological findings.^4,63^ Other studies have reported the slow progress of myelination in Cg, which may explain the relatively early change in MD but no significant increase in FA given the slow microstructural growth.^54^

### Limitations

Due to the challenges in data acquisition and reconstruction, we included 89 out of 135 dMRI scans at GA from 24 to 38 weeks for atlas construction. The number of subjects in each GA week was still limited, although we have performed careful initialization with pairwise registration and shape update to avoid individual bias. A larger sample size is needed to validate the findings, ideally with multi-center data. Also the dMRI data was acquired only in the axial orientation due to the restricted scan time, and thus resolution in the through-slice axis was limited even with super-resolution reconstruction. The proposed atlas generation pipeline together with data collected in multi-orientation would be ideal for further improvement of the atlas. In this work, we approximated the order of WM development based on the turning points in the developmental trajectories of MD, and associated the turning point with the pre-myelination process. However, we were not able to directly measure the exact myelin content in WM tracts, which would require other quantitative markers such as myelin water imaging or magnetic transfer imaging.^64,65^

## Conclusion

In this work, we proposed an FOD-based registration pipeline for fetal brain dMRI atlas generation, combining the pair-wise registration and the kernel-regression technique, which improved registration accuracy compared to the existing approach with tensor-based registration. Using the proposed pipeline, we generated the first spatiotemporal dMRI atlas of Chinese fetal brain from normal fetuses of 24-38 weeks of GA, in both DTI and FOD spaces, providing rich information for deciphering the normal development of fetal brain WM. The atlas can serve as a normative reference of the fetal brain for detection of prenatal abnormalities such as those in fetuses with CHD. The microstructural and volume metric growth of major WM tracts was depicted by the proposed atlas, characterized by non-monotonic changes of dMRI measurements during different stages of gestation, which also suggested the temporal order of WM development in the fetal brain.

## Acknowledgements

The authors want to thank Maria Deprez and Alena Uus from King’s College London for their technical support.

This work was supported by the Ministry of Science and Technology of the People’s Republic of China (2018YFE0114600), the National Natural Science Foundation of China (61801424, 81971606, 82122032), and the Science and Technology Department of Zhejiang Province (202006140, 2022C03057).

## References

1. Thomas JL, Spassky N, Villegas EMP, Goujet-Zalc C, Martinez S, Zalc B. Spatiotemporal development of oligodendrocytes in the embryonic brain. Journal of Neuroscience Research. Feb 15 2000;59(4):471–476. doi:10.1002/(SICI)1097-4547(20000215)59:4<471::AID-JNR1>3.0.CO;2-3

2. Dubois J, Dehaene-Lambertz G, Kulikova S, Poupon C, Hueppi PS, Hertz-Pannier L. The Early development of Brain White Matter: A Review of Imaging Studies in Fetuses, Newborns and Infants. Neuroscience. Sep 12 2014;276:48–71. doi:10.1016/j.neuroscience.2013.12.044

3. Ouyang M, Dubois J, Yu Q, Mukherjee P, Huang H. Delineation of early brain development from fetuses to infants with diffusion MRI and beyond. Neuroimage. Jan 15 2019;185:836–850. doi:10.1016/j.neuroimage.2018.04.017

4. Brody BA, Kinney HC, Kloman AS, Gilles FH. Sequence of Central-Nervous-Sysytem Myelination in Human Infancy. 1. An Autopsy Study of Myelination. Journal of Neuropathology and Experimental Neurology. May 1987;46(3):283–301. doi:10.1097/00005072-198705000-00005

5. Kinney HC, Brody BA, Kloman AS, Gilles FH. Sequence of Central-Nervous-Sysytem Myelination in Human Infancy. 2. Patterns of Myelination in Autopsied Infants. Journal of Neuropathology and Experimental Neurology. May 1988;47(3):217–234. doi:10.1097/00005072-198805000-00003

6. Back SA, Luo NL, Borenstein NS, Volpe JJ, Kinney HC. Arrested oligodendrocyte lineage progression during human cerebral white matter development: Dissociation between the timing of progenitor differentiation and myelinogenesis. Journal of Neuropathology and Experimental Neurology. Feb 2002;61(2):197–211. doi:10.1093/jnen/61.2.197

7. Gilles FHS, W.; Dooling, E. C. Chapter 12 - Mylinated Tracts: Growth Patterns. The Developing Human Brain Growth and Epidemiologic Neuropathology. 1983:117–183:chap 12.

8. Tanaka S, Mito T, Takashima S. Progress of Myelination in the Human Fetal Spinal Nerve Roots, Spinal-cord and Brain-stem with Myelin Basic-Protein Immunohistochemistry. Early Human Development. Mar 17 1995;41(1):49–59. doi:10.1016/0378-3782(94)01608-r

9. Pierpaoli C, Jezzard P, Basser PJ, Barnett A, DiChiro G. Diffusion tensor MR imaging of the human brain. Radiology. Dec 1996;201(3):637–648. doi:10.1148/radiology.201.3.8939209

10. Alexander AL, Lee JE, Lazar M, Field AS. Diffusion tensor imaging of the brain. Neurotherapeutics. Jul 2007;4(3):316–329. doi:10.1016/j.nurt.2007.05.011

11. Tournier JD, Calamante F, Connelly A. Robust determination of the fibre orientation distribution in diffusion MRI: Non-negativity constrained super-resolved spherical deconvolution. Neuroimage. May 1 2007;35(4):1459–1472. doi:10.1016/j.neuroimage.2007.02.016

12. Christiaens D, Cordero-Grande L, Pietsch M, et al. Scattered slice SHARD reconstruction for motion correction in multi-shell diffusion MRI. Neuroimage. Jan 15 2021;225117437. doi:10.1016/j.neuroimage.2020.117437

13. Fogtmann M, Seshamani S, Kroenke C, et al. A Unified Approach to Diffusion Direction Sensitive Slice Registration and 3-D DTI Reconstruction From Moving Fetal Brain Anatomy. Ieee Transactions on Medical Imaging. Feb 2014;33(2):272–289. doi:10.1109/tmi.2013.2284014

14. Kim D-H, Chung S, Vigneron DB, Barkovich AJ, Glenn OA. Diffusion-weighted imaging of the fetal brain in vivo. Magnetic Resonance in Medicine. Jan 2008;59(1):216–220. doi:10.1002/mrm.21459

15. Miller JH, McKinstry RC, Philip JV, Mukherjee P, Neil JJ. Diffusion-tensor MR imaging of normal brain maturation: A guide to structural development and myelination. American Journal of Roentgenology. Mar 2003;180(3):851–859. doi:10.2214/ajr.180.3.1800851

16. Qiu A, Mori S, Miller MI. Diffusion Tensor Imaging for Understanding Brain Development in Early Life. Annual Review of Psychology, Vol 66. 2015 2015;66:853–876. doi:10.1146/annurev-psych-010814-015340

17. Righini A, Bianchini E, Parazzini C, et al. Apparent diffusion coefficient determination in normal fetal brain: A prenatal MR imaging study. American Journal of Neuroradiology. May 2003;24(5):799–804.

18. Bui T, Daire J-L, Chalard F, et al. Microstructural development of human brain assessed in utero by diffusion tensor imaging. Pediatric Radiology. Nov 2006;36(11):1133–1140. doi:10.1007/s00247-006-0266-3

19. Schneider JF, Confort-Gouny S, Le Fur Y, et al. Diffusion-weighted imaging in normal fetal brain maturation. European Radiology. Sep 2007;17(9):2422–2429. doi:10.1007/s00330-007-0634-x

20. Schneider MM, Berman JI, Baumer FM, et al. Normative Apparent Diffusion Coefficient Values in the Developing Fetal Brain. American Journal of Neuroradiology. Oct 2009;30(9): 1799–1803. doi:10.3174/ajnr.A1661

21. Jaimes C, Machado-Rivas F, Afacan O, et al. In vivo characterization of emerging white matter microstructure in the fetal brain in the third trimester. Human Brain Mapping. Aug 15 2020;41(12):3177–3185. doi:10.1002/hbm.25006

22. Hooker JD, Khan MA, Farkas AB, et al. Third-trimester in utero fetal brain diffusion tensor imaging fiber tractography: a prospective longitudinal characterization of normal white matter tract development. Pediatric Radiology. Jun 2020;50(7):973–983. doi:10.1007/s00247-020-04639-8

23. Hoffmann C, Weisz B, Lipitz S, et al. Regional apparent diffusion coefficient values in 3rd trimester fetal brain. Neuroradiology. Jul 2014;56(7):561–567. doi:10.1007/s00234-014-1359-6

24. Korostyshevskaya AM, Prihod’ko IY, Savelov AA, Yarnykh VL. Direct comparison between apparent diffusion coefficient and macromolecular proton fraction as quantitative biomarkers of the human fetal brain maturation. Journal of Magnetic Resonance Imaging. Jul 2019;50(1):52–61. doi:10.1002/jmri.26635

25. Zanin E, Ranjeva J-P, Confort-Gouny S, et al. White matter maturation of normal human fetal brain. An in vivo diffusion tensor tractography study. Brain and Behavior. Nov 2011;1(2):95–108. doi:10.1002/brb3.17

26. Wilson S, Pietsch M, Cordero-Grande L, et al. Development of human white matter pathways in utero over the second and third trimester. Proceedings of the National Academy of Sciences of the United States of America. May 18 2021;118(20) e2023598118. doi:10.1073/pnas.2023598118

27. Machado-Rivas F, Afacan O, Khan S, et al. Spatiotemporal changes in diffusivity and anisotropy in fetal brain tractography. Human Brain Mapping. 2021;doi:10.1002/hbm.25653

28. Khan S, Vasung L, Marami B, et al. Fetal brain growth portrayed by a spatiotemporal diffusion tensor MRI atlas computed from in utero images. Neuroimage. Jan 15 2019;185:593–608. doi:10.1016/j.neuroimage.2018.08.030

29. Lou Y, Zhao L, Yu S, et al. Brain asymmetry differences between Chinese and Caucasian populations: a surface-based morphometric comparison study. Brain Imaging and Behavior. Dec 2020;14(6):2323–2332. doi:10.1007/s11682-019-00184-7

30. Tang Y, Hojatkashani C, Dinov ID, et al. The construction of a Chinese MRI brain atlas: A morphometric comparison study between Chinese and Caucasian cohorts. Neuroimage. May 15 2010;51(1):33–41. doi:10.1016/j.neuroimage.2010.01.111

31. Tang Y, Zhao L, Lou Y, et al. Brain structure differences between Chinese and Caucasian cohorts: A comprehensive morphometry study. Human Brain Mapping. May 2018;39(5):2147–2155. doi:10.1002/hbm.23994

32. Zhao T, Liao X, Fonov VS, et al. Unbiased age-specific structural brain atlases for Chinese pediatric population. Neuroimage. Apr 1 2019;189:55–70. doi:10.1016/j.neuroimage.2019.01.006

33. Chansangpetch S, Huang G, Coh P, et al. Differences in Optic Nerve Head, Retinal Nerve Fiber Layer, and Ganglion Cell Complex Parameters Between Caucasian and Chinese Subjects. Journal of Glaucoma. Apr 2018;27(4):350–356. doi:10.1097/ijg.0000000000000889

34. Bai J, Abdul-Rahman MF, Rifkin-Graboi A, et al. Population Differences in Brain Morphology and Microstructure among Chinese, Malay, and Indian Neonates. Plos One. Oct 24 2012;7(10) e47816. doi:10.1371/journal.pone.0047816

35. Gynecology ISoUiO. Cardiac screening examination of the fetus: guidelines for performing the ‘basic’ and ‘extended basic’ cardiac scan. Ultrasound in Obstetrics & Gynecology. Jan 2006;27(1):107–113. doi:10.1002/uog.2677

36. Tournier JD, Smith R, Raffelt D, et al. MRtrix3: A fast, flexible and open software framework for medical image processing and visualisation. Neuroimage. Nov 15 2019;202116137. doi:10.1016/j.neuroimage.2019.116137

37. Deprez M, Price A, Christiaens D, et al. Higher Order Spherical Harmonics Reconstruction of Fetal Diffusion MRI With Intensity Correction. Ieee Transactions on Medical Imaging. Apr 2020;39(4): 1104–1113. doi:10.1109/tmi.2019.2943565

38. Kuklisova-Murgasova M, Quaghebeur G, Rutherford MA, Hajnal JV, Schnabel JA. Reconstruction of fetal brain MRI with intensity matching and complete outlier removal. Medical Image Analysis. Dec 2012;16(8):1550–1564. doi:10.1016/j.media.2012.07.004

39. Raffelt DA, Tournier JD, Smith RE, et al. Investigating white matter fibre density and morphology using fixel-based analysis. Neuroimage. Jan 1 2017;144:58–73. doi:10.1016/j.neuroimage.2016.09.029

40. Serag A, Aljabar P, Ball G, et al. Construction of a consistent high-definition spatio-temporal atlas of the developing brain using adaptive kernel regression. Neuroimage. Feb 1 2012;59(3):2255–2265. doi:10.1016/j.neuroimage.2011.09.062

41. Davis BC, Fletcher PT, Bullitt E, Joshi S. Population Shape Regression from Random Design Data. International Journal of Computer Vision. Nov 2010;90(2):255–266. doi:10.1007/s11263-010-0367-1

42. Serag A, Aljabar P, Ball G, et al. Construction of a consistent high-definition spatio-temporal atlas of the developing brain using adaptive kernel regression (vol 59, pg 2255, 2012). Neuroimage. Nov 1 2012;63(2):998–998. doi:10.1016/j.neuroimage.2012.01.086

43. Raffelt D, Tournier JD, Fripp J, Crozier S, Connelly A, Salvado O. Symmetric diffeomorphic registration of fibre orientation distributions. Neuroimage. Jun 1 2011;56(3):1171–1180. doi:10.1016/j.neuroimage.2011.02.014

44. Alexander DC, Pierpaoli C, Basser PJ, Gee JC. Spatial transformations of diffusion tensor magnetic resonance images. Ieee Transactions on Medical Imaging. Nov 2001;20(11):1131–1139. doi:10.1109/42.963816

45. Tournier JDC, Fernando; Connelly, Alan. Improved probabilistic streamlines tractography by 2nd order integration over fibre orientation distributions. 2010:

46. Wang J, Aydogan DB, Varma R, Toga AW, Shi Y. Topographic Regularity for Tract Filtering in Brain Connectivity. Information Processing in Medical Imaging (Ipmi 2017). 2017 2017; 10265:263–274. doi:10.1007/978-3-319-59050-9_21

47. Yin XY, Goudriaan J, Lantinga EA, Vos J, Spiertz HJ. A flexible sigmoid function of determinate growth (vol 91, pg 361, 2003). Annals of Botany. May 2003;91(6):753–753.

48. Raffelt DA, Smith RE, Ridgway GR, et al. Connectivity-based fixel enhancement: Whole-brain statistical analysis of diffusion MRI measures in the presence of crossing fibres. Neuroimage. Aug 15 2015;117:40–55. doi:10.1016/j.neuroimage.2015.05.039

49. Smith SM, Nichols TE. Threshold-free cluster enhancement: Addressing problems of smoothing, threshold dependence and localisation in cluster inference. Neuroimage. Jan 1 2009;44(1):83–98. doi:10.1016/j.neuroimage.2008.03.061

50. Hagmann C, Singer J, Latal B, Knirsch W, Makki M. Regional Microstructural and Volumetric Magnetic Resonance Imaging (MRI) Abnormalities in the Corpus Callosum of Neonates With Congenital Heart Defect Undergoing Cardiac Surgery. Journal of Child Neurology. Mar 2016;31(3):300–308. doi:10.1177/0883073815591214

51. Perez-Cruz M, Gomez O, Gibert M, et al. Corpus callosum size by neurosonography in fetuses with congenital heart defect and relationship with expected pattern of brain oxygen supply. Ultrasound in obstetrics & gynecology : the official journal of the International Society of Ultrasound in Obstetrics and Gynecology. 2021-May-16 2021;doi:10.1002/uog.23684

52. Oishi K, Mori S, Donohue PK, et al. Multi-contrast human neonatal brain atlas: Application to normal neonate development analysis. Neuroimage. May 1 2011;56(1):8–20. doi:10.1016/j.neuroimage.2011.01.051

53. Huang H, Zhang J, Wakana S, et al. White and gray matter development in human fetal, newborn and pediatric brains. Neuroimage. Oct 15 2006;33(1):27–38. doi:10.1016/j.neuroimage.2006.06.009

54. Kinney HCV, Joseph J. Chapter 8 - Myelination Events. In: Volpe JJI, Terrie E.; Darras, Basil T.; De Vries, Linda S.; Du Plessis, Adré J.; Neil, Jeffrey J.; Perlman, Jeffrey M., ed. Volpe’s Neurology of the Newborn (Sixth Edition). 6 ed. 2018:176–188:chap 8.

55. Dubois J, Dehaene-Lambertz G, Perrin M, et al. Asynchrony of the early maturation of white matter bundles in healthy infants: Quantitative landmarks revealed noninvasively by diffusion tensor imaging. Human Brain Mapping. Jan 2008;29(1):14–27. doi:10.1002/hbm.20363

56. Counsell SJ, Maalouf EF, Fletcher AM, et al. MR imaging assessment of myelination in the very preterm brain. American Journal of Neuroradiology. May 2002;23(5):872–881.

57. McKinstry RC, Mathur A, Miller JH, et al. Radial organization of developing preterm human cerebral cortex revealed by non-invasive water diffusion anisotropy MRI. Cerebral Cortex. Dec 2002;12(12):1237–1243. doi:10.1093/cercor/12.12.1237

58. Takahashi E, Folkerth RD, Galaburda AM, Grant PE. Emerging Cerebral Connectivity in the Human Fetal Brain: An MR Tractography Study. Cerebral Cortex. Feb 2012;22(2):455–464. doi:10.1093/cercor/bhr126

59. Licht DJ, Shera DM, Clancy RR, et al. Brain maturation is delayed in infants with complex congenital heart defects. Journal of Thoracic and Cardiovascular Surgery. Mar 2009;137(3):529–537. doi:10.1016/j.jtcvs.2008.10.025

60. Clouchoux C, du Plessis AJ, Bouyssi-Kobar M, et al. Delayed Cortical Development in Fetuses with Complex CHD. Cerebral Cortex. Dec 2013;23(12):2932–2943. doi:10.1093/cercor/bhs281

61. Jaimes C, Rofeberg V, Stopp C, et al. Association of Isolated CHD with Fetal Brain Maturation. American Journal of Neuroradiology. Aug 2020;41(8):1525–1531. doi:10.3174/ajnr.A6635

62. Milos R-I, Jovanov-Milosevic N, Mitter C, et al. Developmental dynamics of the periventricular parietal crossroads of growing cortical pathways in the fetal brain - In vivo fetal MRI with histological correlation. Neuroimage. Apr 15 2020;210116553. doi:10.1016/j.neuroimage.2020.116553

63. Huang H, Vasung L. Gaining insight of fetal brain development with diffusion MRI and histology. International Journal of Developmental Neuroscience. Feb 2014;32:11–22. doi:10.1016/j.ijdevneu.2013.06.005

64. Nossin-Manor R, Card D, Morris D, et al. Quantitative MRI in the very preterm brain: Assessing tissue organization and myelination using magnetization transfer, diffusion tensor and T-1 imaging. Neuroimage. Jan 1 2013;64:505–516. doi:10.1016/j.neuroimage.2012.08.086

65. Weber AM, Zhang Y, Kames C, Rauscher A. Myelin water imaging and R-2* mapping in neonates: Investigating R-2* dependence on myelin and fibre orientation in whole brain white matter. Nmr in Biomedicine. Mar 2020;33(3) e4222. doi:10.1002/nbm.4222

