## supplementary for "Deciphering the developmental order and microstructural patterns of early white matter pathways in a diffusion MRI based fetal brain atlas"

### Supplementary Material

#### The fetal brain dMRI atlas

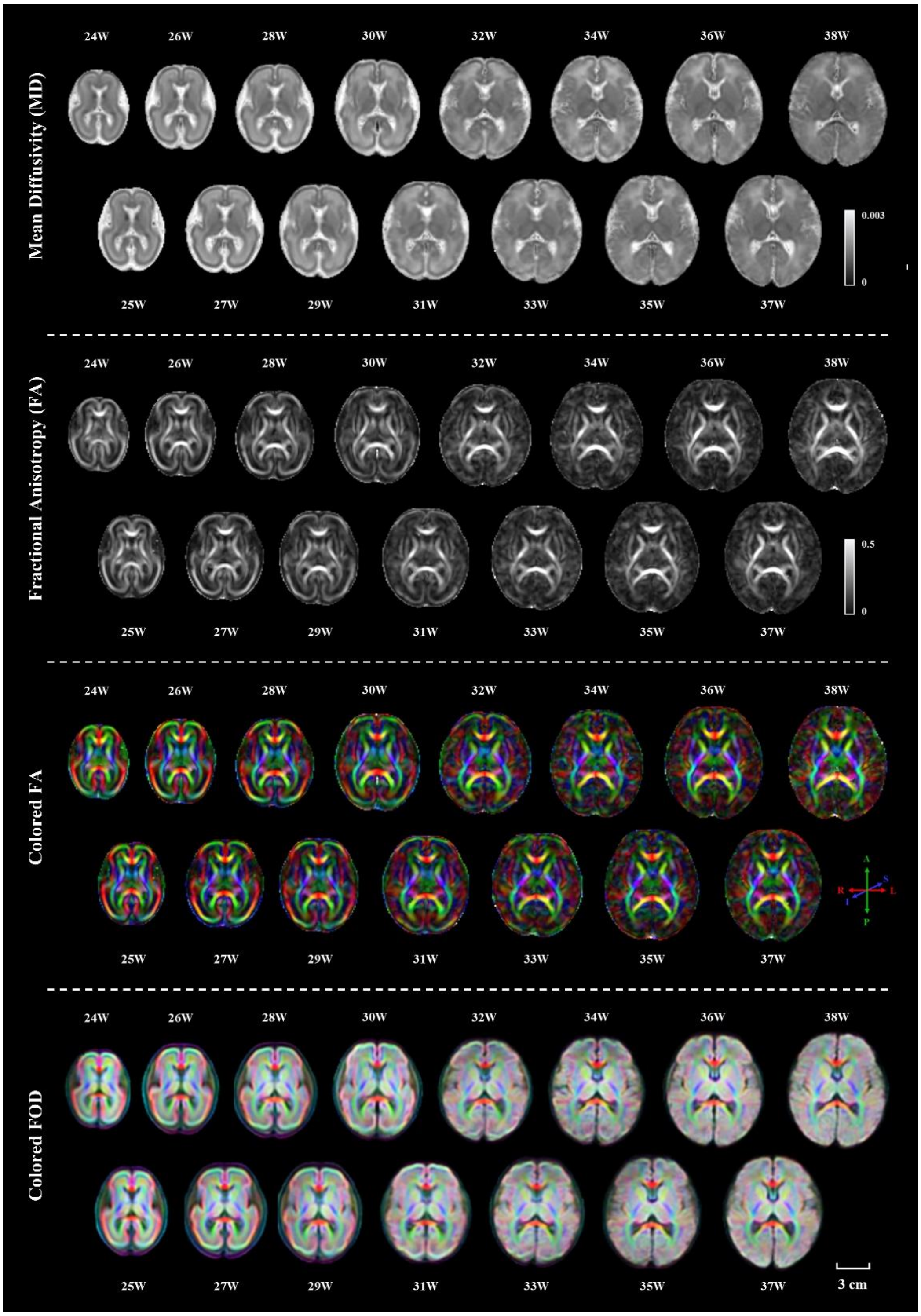


**Supplementary Figure 1  Multi-modal fetal brain dMRI atlas from 24-38 weeks including both FOD and DTI metrics.**

#### ROI definitions for fiber tracking

Tractography in the atlas space was performed in MRtrix3 using the probabilistic algorithm iFOD2.[1] For each WM tract, at least one seed ROI was manually delineated, and segmentation-based or manually drawn ROIs were used as inclusion or exclusion regions. We referred to the WM segmentations in JHU single subject neonatal atlas for the manual ROI delineation.[2] To obtain segmentation-based ROIs, we non-linearly registered the mean diffusivity (MD) templates to the CRL fetal brain T2 weighted atlas of corresponding gestational ages (GAs), and T2 segmentations of the CRL atlas were transformed to our atlas space using the inverse non-linear warps. The manually delineated ROIs are displayed in Figure S2 and the detailed criterion for ROI delineation for each tract are described below.

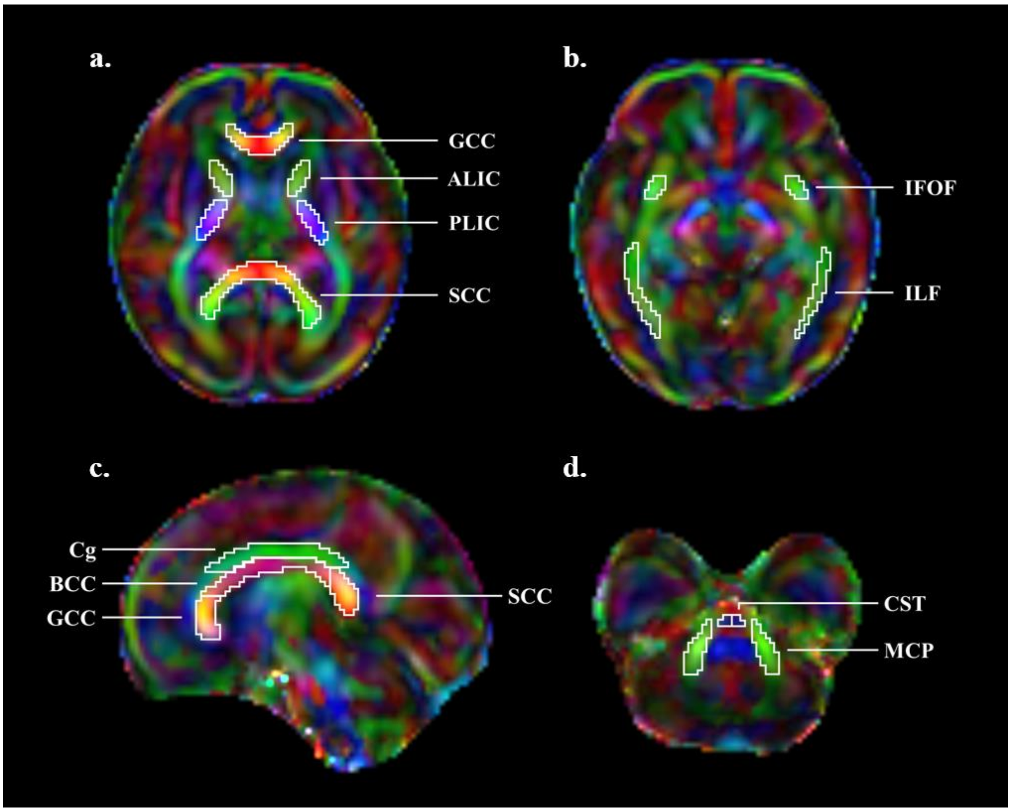


**Supplementary Figure 2 Manually delineated regions of interest (ROIs).** ROIs including the genu, body and splenium of corpus callosum (GCC, BCC and SCC), and bilateral anterior limb of internal capsule (ALIC), posterior limb of internal capsule (PLIC), Cingulum bundle (Cg), inferior fontal occipital fasciculus (IFOF), inferior longitudinal fasciculus (ILF), middle cerebellar peduncle (MCP) and the cortico-spinal tract (CST) were manually delineated. The ROIs were used as seed regions for fiber tractography, and as the regions for ROI-based comparison between normal and congenital heart disease (CHD) fetal brains.

#### Cortico-spinal tract (CST)

A seed ROI was defined in the CST at the brainstem level, which can be visualized as blue areas in the inferior part of the brain on the colored FA maps (Figure S2d). Left and right posterior limb of the internal capsule (PLIC) were used as waypoints (Figure S2a). ROIs were manually drawn on axial slices.

#### Inferior fronto-occipital fasciculus (IFOF)

The seed region was drawn on axial slices in the IFOF close to the external capsule (green ROIs on colored FA maps below the external capsules, Figure S2b). The frontal lobe and occipital lobes obtained from the T2 segmentation were set as included regions.

#### Inferior longitudinal fasciculus (ILF)

ILF was traced using the T2 segmentations of occipital lobes as the seed regions and the temporal lobes as inclusion areas. A waypoint ROI manually delineated in ILF (Figure S2b) was also defined as an inclusion area.

#### Cingulum bungle (Cg)

A single ROI was delineated on 1- 3 sagittal slices around the midline as the seed region for Cg, covering a slim green area superior to CC (Figure S2c). In brain regions superior to Cg, there are complex cortico-cortical connection fibers, and some of them run in the same direction as Cg.[3] Because of the limited spatial resolution, the boundaries between the cingulum bundle and these fiber structures superior are blurred, especially in GAs earlier than 31W due to strong partial volume effect. Therefore, exclusion areas superior to the cingulum in cortical regions were manually delineated to reduce false positives.

#### Corpus callosum (CC)

As shown in Figure S2c, the genu of CC (GCC) was drawn in the anterior part of CC where FA values were relatively high and the splenium of CC (SCC) was drawn in a similar manner in the posterior part of CC. The middle part where FA was relatively low was segmented as the body of CC (BCC). We used these ROIs as seed regions and removed false positive streamlines extending to the deep gray matter regions of the brain using axial or coronal slices as exclusion ROIs.

##### Middle cerebellar peduncle (MCP)

For both left and right MCP, an area along this tract in the brainstem with relatively high FA was drawn on an axial slice, which is a green area in the colored FA maps (Figure S2d). We used the left MCP as seed region and included right MCP.[4] An axial slice at the upper border of MCP was defined as the exclusion ROI.

#### Fixel-based analysis (FBA)

We used the FBA pipeline developed by MRtrix3 to study fiber density (FD), fiber-bundle cross-section (FC), and fiber density and cross-section (FDC) values in WM fiber bundles. In the pipeline, FD values were directly calculated from the subject FODs as the integration of FOD amplitude across all lobes.[5] To compute FC, a population-based template needs to be built and the FC value is calculated from the subject-to-template warps, simply as the overall volume change (Jacobian determinant) factoring out the change along the direction of the fixel. FDC is computed by multiplying FD and FC. To perform fixel-based statistical analysis, FD, FC, and FDC values in the subject fixel space were mapped to the population template space. Since the fetal population in this study contained fetuses from 24W to 38W of gestation, there are significant anatomical differences between subjects. The resulting population-based template only reserved the most commonly shared structural characteristics (the most common fixels) of these subjects and omitted some detailed information (cross-fibers at later stage, Figure S3). Therefore, and the statistical analysis was performed for the common "principle fixels". To set a uniform standard, the sampling of the FBA-based parameters in each tract was also performed in the common template space. To convert fixel-based values to voxel-wise images, we computed the sum of these values in each voxels to represent the overall characteristics of microstructure. On the other hand, the calculation of FD does not need a template so we can obtain both the original FD in subject spaces and averaged FD in the template space. The FD values sampled from subject spaces were displayed in Figure S4, which were slightly different from the main results (Figure 5a) obtained in the template space. In the subject fixel space, the sum of FD was higher than that in the template space due to the inclusion of complex cross-fiber fixels (Figure S3, left). Compared to FD values sampled from the template space, FD in the subject space decreased slower with GA, and in IFOF, ILF, SCC, and GCC, the FD value showed a slight increase in the late third trimester (Figure S4) indicating the increase of complexity at late gestation.

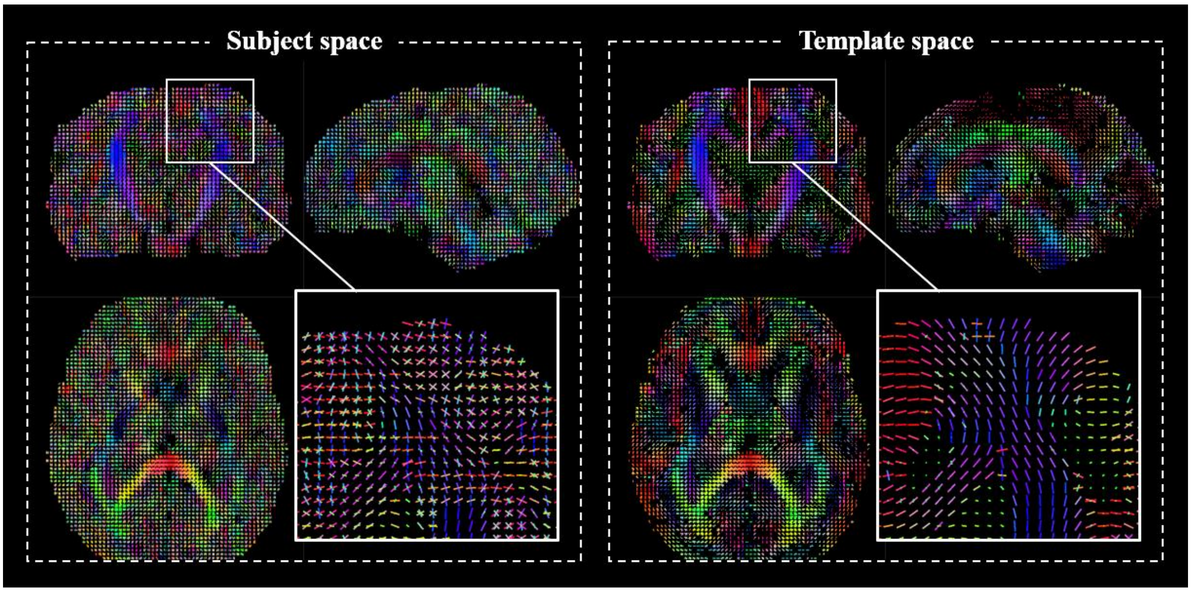


**Supplementary Figure 3 Comparison of fixels in the subject space and the template space.** Fixels of a 38W fetal brain (left panel) were mapped to the population-based template space on the left panel.

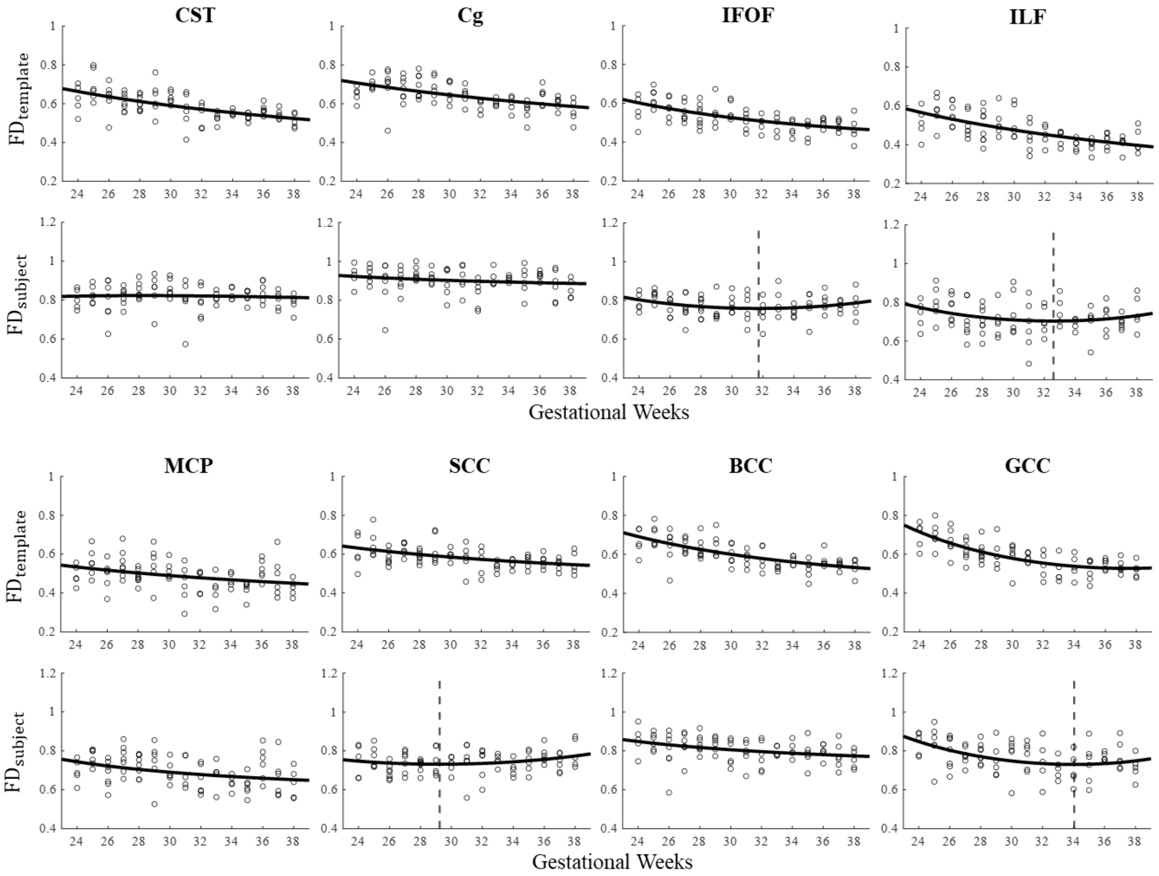


**Supplementary Figure 4 Developmental trajectories of FD in WM tracts sampled from the template space (first row) and subject spaces (second row).**

#### Comparison with the CRL fetal brain DTI atlas

We put the generated Chinese fetal brain dMRI atlas and the CRL fetal brain DTI atlas (http://crl.med.harvard.edu/research/fetal_brain_atlas/) together in Figure S5 for visual comparison. As displayed, the two atlases revealed similar developmental patterns of the fetal brain, as the colored FA maps showed the emerging complex WM tracts and cortical microstructures with GA. However, the CRL DTI atlas has poorer in-plane resolution compared to our atlas. The cortical details displayed by our colored FA maps are not visible in the CRL atlas (Figure S5, right panel).

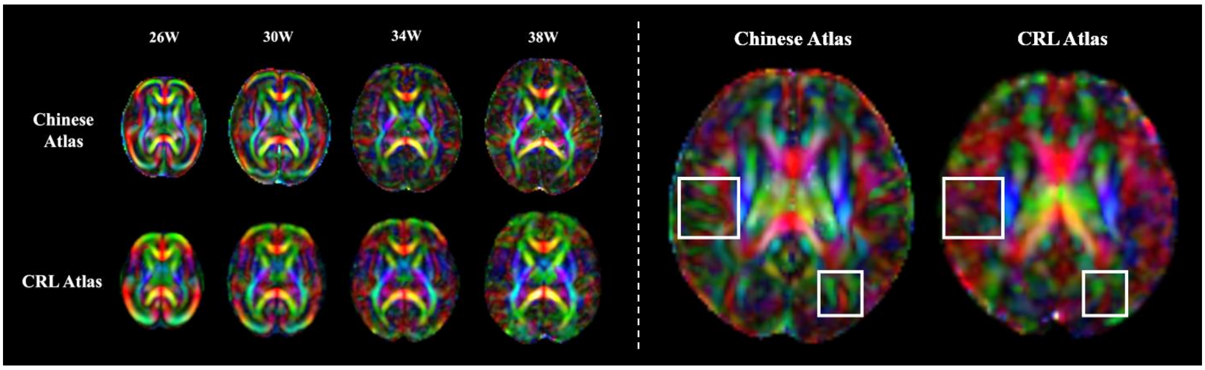


**Supplementary Figure 5  Comparison between the Chinese fetal brain dMRI atlas and the CRL atlas.** The CRL atlas has poorer in-plane spatial resolution compared to our atlas. The dMRI atlas that we generated displays sharper structural details in cortical regions, e.g., in the 36 week template.

To quantitatively compare the developmental patterns between Chinese and Caucasian fetal brain atlas, we evaluated FA and MD values of the 8 major WM tracts in these two atlas. Since the CRL did not provide the individual subject data but only the final DTI atlas, the comparison can only be performed between population-averaged templates. Each MD and FA maps of CRL atlas were non-linearly registered to our atlas of corresponding GAs thorough multi-channel diffeomorphic registration using ANTS.[6] Then we sampled mean FA and MD value in both atlases using the same sets of WM ROIs (Figure S2) from each GA. The results are shown in Figure S6.

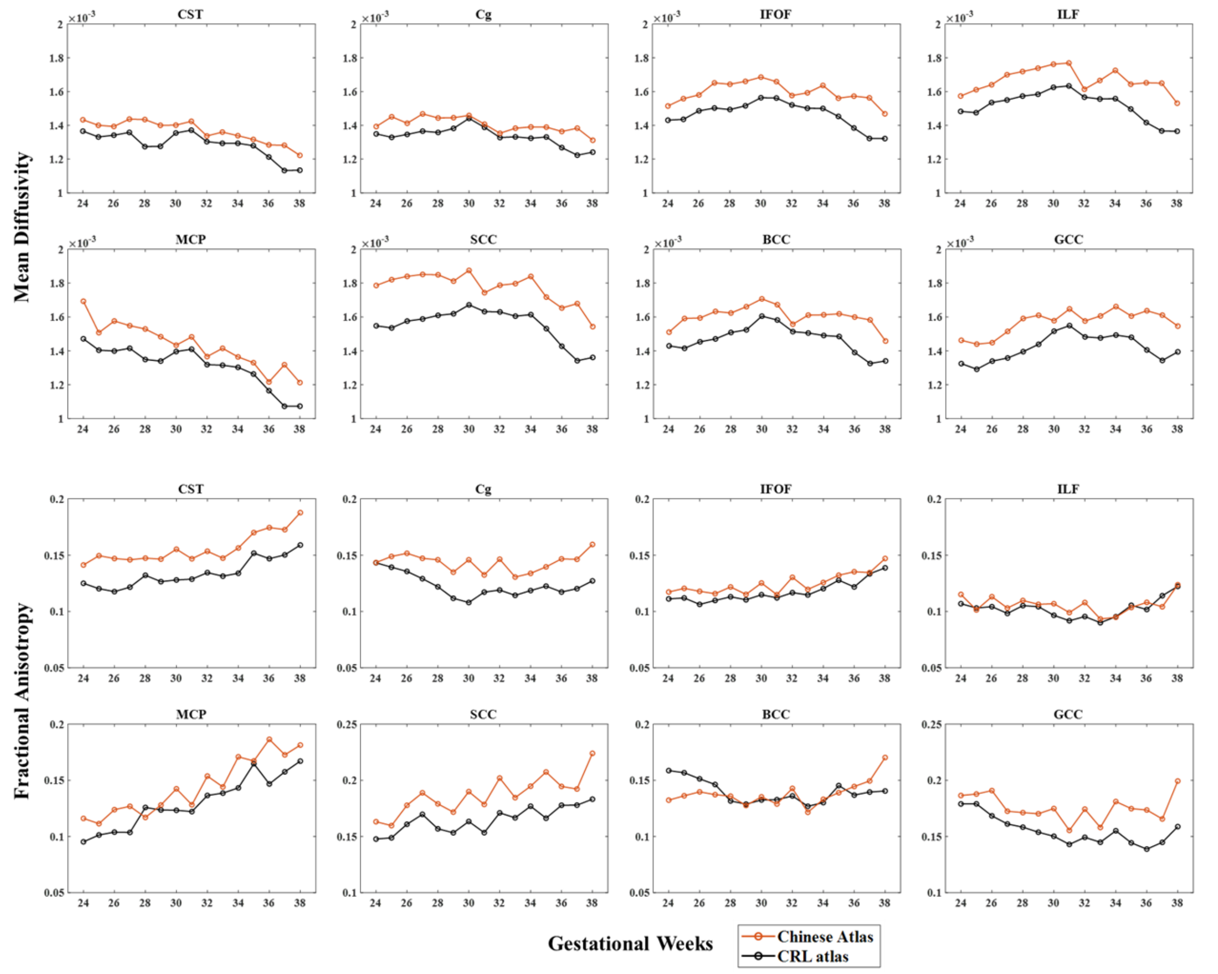


**Supplementary Figure 6 Comparison of DTI-derived measurements in WM ROIs between the Chinese atlas and the CRL atlas.** Similar trends of FA and MD were found in these two atlases but the absolute values were different.

We found similar developmental trends between the Chinese and Caucasian fetal brain, but the MD and FA values in these two atlases were different. It is hard to determine if these differences are biology, because the different imaging protocols and reconstruction methods can largely influence the dMRI data. The CRL dMRI data were acquired from multiple stacks including the axial, coronal and sagittal views, but the data had lower in-plane resolution (2mm) and poor angular resolution (12 gradient directions) compared to our data. Moreover, they used the weighted average of the unregistered subject image in tensor space to initialize the atlas rather than the pair-wise registration in FOD space in our pipeline, which may introduce blurring to the final atlas.

To examine whether the reconstruction methods contribute to variations in DTI-derived metrics, we managed to obtain axial, coronal, and sagittal stacks from 5 normal fetuses (GAs were 26W, 28W, 30W, 32W, and 36W). Images were scanned with the same acquisition protocol and went through the same preprocessing steps. We performed two types of reconstructions for each subject, one using all three stacks and the other using one axial stack only. Results in Figure S7 showed the FA and MD values were influenced by different reconstruction methods and the relative differences varied among subjects in different WM structures, suggesting the importance for fixing the acquisition and reconstruction for investigation of biological difference between atlases.

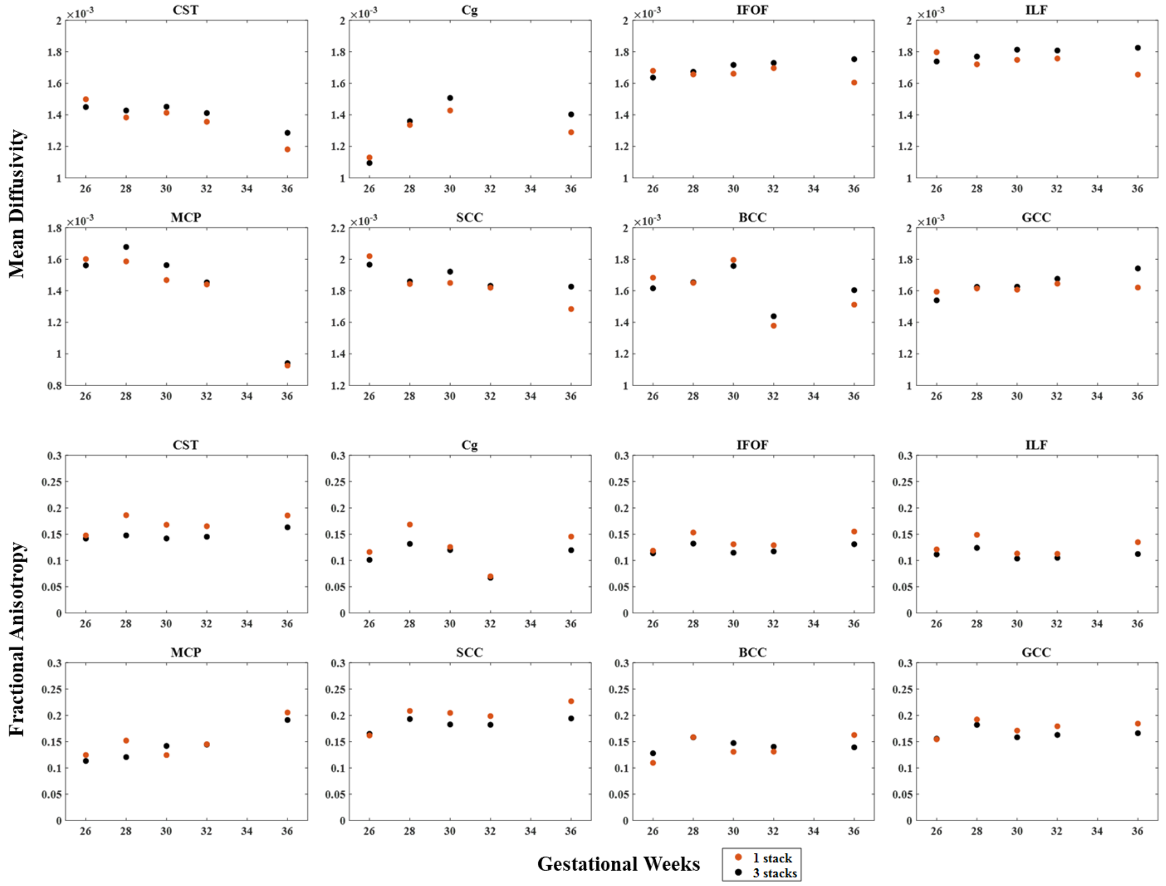


**Supplementary Figure 7 Comparison of DTI-derived measurements in WM tracts between data reconstructed from one axial stack and orthogonal stacks in three directions.** The FA and MD values were influenced by different reconstruction methods and the relative differences varied among subjects in different WM structures.

### Reference

1. Tournier JD, Smith R, Raffelt D, et al. MRtrix3: A fast, flexible and open software framework for medical image processing and visualisation. *Neuroimage*. Nov 15 2019;202116137. doi:10.1016/j.neuroimage.2019.116137

2. Oishi K, Mori S, Donohue PK, et al. Multi-contrast human neonatal brain atlas: Application to normal neonate development analysis. *Neuroimage*. May 1 2011;56(1):8-20. doi:10.1016/j.neuroimage.2011.01.051

3. Takahashi E, Folkerth RD, Galaburda AM, Grant PE. Emerging Cerebral Connectivity

in the Human Fetal Brain: An MR Tractography Study. *Cerebral Cortex*. Feb 2012;22(2):455-464. doi:10.1093/cercor/bhr126

4. Leitner Y, Travis KE, Ben-Shachar M, Yeom KW, Feldman HM. Tract Profiles of the Cerebellar White Matter Pathways in Children and Adolescents. *Cerebellum*. Dec 2015;14(6):613-623. doi:10.1007/s12311-015-0652-1

5. Raffelt DA, Tournier JD, Smith RE, et al. Investigating white matter fibre density and morphology using fixel-based analysis. *Neuroimage*. Jan 1 2017;144:58-73. doi:10.1016/j.neuroimage.2016.09.029

6. Avants BB, Epstein CL, Grossman M, Gee JC. Symmetric diffeomorphic image registration with cross-correlation: Evaluating automated labeling of elderly and neurodegenerative brain. *Medical Image Analysis*. Feb 2008;12(1):26-41. doi:10.1016/j.media.2007.06.004
